# Maternal H3K36 and H3K27 HMTs protect germline immortality via regulation of the transcription factor LIN-15B

**DOI:** 10.1101/2022.03.17.484692

**Authors:** Chad Cockrum, Susan Strome

## Abstract

Maternally synthesized products play critical roles in development of offspring. A premier example is the *C. elegans* H3K36 methyltransferase MES-4, which is essential for germline survival and development in offspring. How maternal MES-4 protects germline immortality is not well understood, but its role in H3K36 methylation hinted that it may regulate gene expression in Primordial Germ Cells (PGCs). We tested this hypothesis by profiling transcripts from single pairs of PGCs dissected from wild-type and *mes-4* mutant (lacking maternal and zygotic MES-4) newly hatched larvae. We found that *mes-4* PGCs display normal turn-on of most germline genes and normal repression of somatic genes, but dramatically up-regulate hundreds of genes on the X chromosome. We demonstrated that X mis-expression is the cause of germline death by generating and analyzing *mes-4* mutants that inherited different endowments of X chromosome(s). Intriguingly, removal of the THAP transcription factor LIN-15B from *mes-4* mutants reduced X mis-expression and prevented germline death. *lin-15B* is X-linked and mis-expressed in *mes-4* PGCs, identifying it as a critical target for MES-4 repression. The above findings extend to the H3K27 methyltransferase MES-2/3/6, the *C. elegans* version of Polycomb Repressive Complex 2. We propose that maternal MES-4 and PRC2 cooperate to protect germline survival by preventing synthesis of germline-toxic products encoded by genes on the X chromosome, including the key transcription factor LIN-15B.

## INTRODUCTION

Many critical events during early development are orchestrated by maternally synthesized gene products. Mutations in genes that encode such products in the mother can cause ‘maternal-effect’ phenotypes in offspring. These phenotypes are usually severe developmental defects. Maternal-effect lethal genes, which cause maternal-effect death of offspring, encode products that guide crucial events in early embryo development, such as pattern formation and embryonic genome activation (e.g. the PAR proteins in *C. elegans*, BICOID in *Drosophila*, and Mater in mouse) (Nusslein-Volhard et al. 1987; Tong et al., 2000; Kemphues et al. 1988). Maternal-effect sterile genes encode products needed for fertility of the offspring. A few genes in this category encode proteins in germ granules (e.g. PGL-1 in *C. elegans* and VASA in *Drosophila*) (Nusslein-Volhard et al. 1987; Rongo and Lehmann, 1996; Kawasaki et al., 1998).

Another fascinating set of genes in this category encode chromatin regulators, which are the focus of this paper. The *C. elegans* MES proteins were identified in genetic screens for maternal-effect sterile mutants, hence their name (MES for Maternal-Effect Sterile) (Capowski et al., 1991). MES-2, MES-3, and MES-6 assemble into a trimeric complex that is the *C. elegans* version of Polycomb Repressive Complex 2 (PRC2) (Xu et al., 2001; Bender et al., 2004). PRC2 is a histone methyltransferase (HMT) that methylates Lys 27 on histone H3 (H3K27me) to repress genes that are packaged by those methylated nucleosomes (Ketel et al., 2005; Margueron and Reinberg, 2011; Pengelly et al., 2013). MES-4 is an HMT that methylates Lys 36 on H3 (H3K36me), which marks actively transcribing genes and has context-dependent roles in transcriptional regulation (Bender et al., 2006; Furuhashi et al., 2010; Rechtsteiner et al., 2010, Kreher et al., 2018). Although PRC2 and MES-4 catalyze opposing flavors of histone marking, the loss of either causes nearly identical mutant phenotypes (Capowski et al., 1991). Worms that inherit a maternal load of gene product but cannot synthesize zygotic product (referred to as *mes* M+Z- mutants) are fertile. Worms that do not inherit a maternal load or produce zygotic gene product (*mes* M-Z- mutants) are sterile due to death of nascent germ cells in early- to mid-stage larvae. In *mes* M-Z+ mutants, zygotically synthesized product does not rescue fertility, highlighting the critical importance of maternal product. PRC2’s roles in transcriptional repression and development have been intensively studied and are well defined across species, including roles in *C. elegans* germline development (Bender et al., 2004, Patel et al., 2012; Gaydos et al., 2014; Kaneshiro et al., 2019; Delaney et al., 2019). In contrast, how MES-4 ensures the survival of nascent germ cells is unknown and particularly puzzling.

One possibility for MES-4 function is that maternal MES-4 promotes expression in offspring of genes required for germline development. Support for this comes from analyses of mutants that ectopically express germline genes in their soma (e.g. *mep-1, lin-15B, lin-35,* and *spr-5; met-2* mutants), and as a result, have developmental defects (Unhavaithaya et al., 2002; Wang et al., 2005; Cui et al., 2006; Petrella et al., 2011; Wu et al., 2012; Carpenter et al., 2021). Concomitant loss of MES-4 from these mutants prevents ectopic expression of germline genes and restores worm health (Wang et al., 2005; Cui et al., 2006; Petrella et al., 2011; Wu et al., 2012; Carpenter et al., 2021). In wild-type early embryos, MES-4 and methylated H3K36 associate with genes that were transcribed in the maternal germline, regardless of whether they are transcribed in embryos (Furuhashi et al., 2010; Rechtsteiner et al., 2010). Focusing on H3K36me3, genetic tests showed that MES-4 strictly maintains pre-existing patterns of H3K36me3 and is unable to catalyze *de novo* H3K36me3 marking of genes (Furuhashi et al., 2010); the other H3K36 HMT in *C. elegans*, MET-1, like H3K36 HMTs in other systems, catalyzes *de novo* H3K36me3 on genes in response to transcriptional turn-on (Kizer et al., 2005; Furuhashi et al., 2010; Kreher et al., 2018). Taken together, these findings suggested the appealing model that in embryos maternal MES-4 maintains H3K36me3 marking of germline-expressed genes and in that way transmits an epigenetic ‘memory of germline’, a developmental blueprint, to the primordial germ cells (PGCs) of offspring.

Two findings challenge the model that MES-4 somehow promotes expression of germline genes. First, among *mes-4* M-Z-mutants, hermaphrodites (with 2 X chromosomes) are always sterile, while males (with 1 X chromosome) can be fertile (Garvin et al., 1998). This suggested that the dosage of X-linked genes matters for the Mes-4 mutant phenotype. Second, profiling transcripts in the gonads of fertile *mes-4* M+Z- mutant hermaphrodites revealed that the most dramatic effect of losing zygotic MES-4 is up-regulation of genes on the X (Bender et al., 2006; Gaydos et al., 2012). Notably, the X chromosomes are normally kept globally repressed during all stages of germline development in males and during most stages of germline development in hermaphrodites (Kelly et al., 2002; Reinke et al., 2004; Wang et al., 2009, Arico et al., 2011; Strome et al., 2014; Tzur et al., 2018), and likely as a consequence, most germline-expressed genes are located on the autosomes. These findings focused attention on the X chromosome and raised the question – what role does maternal MES-4 serve to ensure that PGCs survive and develop into a full and healthy germline?

To investigate the role of MES-4 in PGCs, the cells that critically rely on maternal MES-4 for survival, and to formally test the model that MES-4 promotes expression of germline genes, we performed transcript profiling in dissected single pairs of PGCs from wild-type versus *mes-4* M- Z- mutant larvae. We asked if absence of maternal MES-4 causes PGCs to 1) fail to turn on germline genes, 2) inappropriately turn on somatic genes, and/or 3) inappropriately turn on X-linked genes. We found that in *mes-4* PGCs most germline genes were turned on normally, thus disproving the model that the major role of MES-4 is to promote expression of germline genes in PGCs. Most somatic genes were kept off, arguing that MES-4 does not protect the germline by opposing somatic development. The most dramatic impact to the transcriptome in *mes-4* PGCs was up-regulation of hundreds of X-linked genes, many of which are part of an oogenesis program. Our genetic analysis of *mes-4* mutants with different X-chromosome endowments from the oocyte and sperm demonstrated that up-regulation of X-linked genes is the cause of death of nascent germ cells in *mes-4* M-Z-mutant larvae. We identified the transcription factor LIN-15B, an X-linked gene up-regulated in *mes-4* mutants, as a major cause of X mis-expression and germline death in *mes-4* mutants. Performing similar tests of PRC2 (*mes-3*) M-Z-mutant larvae revealed that their PGCs up-regulate many X-linked genes in common with *mes-4* PGCs, and that removal of LIN-15B restores the health of their germline, as it does for *mes-4* mutants. This study revealed that maternal MES-4 and PRC2 cooperate to ensure germline survival and health in offspring by preventing mis-expression of genes on the X chromosome, and that both operate through a key X-encoded transcription factor.

## RESULTS

### MES-4 is not required for PGCs to launch a germline program

MES-4 propagates an epigenetic ‘memory’ of a germline gene expression program during embryogenesis by maintaining H3K36me3 on genes that were previously transcribed in parental germlines (Rechtsteiner et al., 2010, Furuhasi et al., 2010; Kreher et al., 2018). A popular model predicts that delivery of this memory to offspring PGCs instructs them to launch a gene expression program that promotes germline proliferation and development. To test this model, we performed RNA-sequencing to determine whether PGCs from *mes-4* M-Z-(<u>M</u>aternal MES-4 minus and <u>Z</u>ygotic MES-4 minus) mutant larvae, which completely lack MES-4, fail to turn on a germline program (Figure 1A). We developed a hand-dissection strategy that enables us to isolate in <30 minutes single sets of 2 sister PGCs, marked by a specifically and highly expressed germline marker (GLH-1::GFP), from wild-type or *mes-4* M-Z-mutant larvae for RNA-seq library preparation. We performed differential expression analysis to identify genes that are significantly down-regulated (DOWN) or up-regulated (UP) in *mes-4* mutant PGCs compared to wild-type (wt) PGCs. Our analysis identified 176 DOWN genes and 450 UP genes (Figure 1B-D, Figure 2—figure supplement 2A,B).

**Figure 1:**
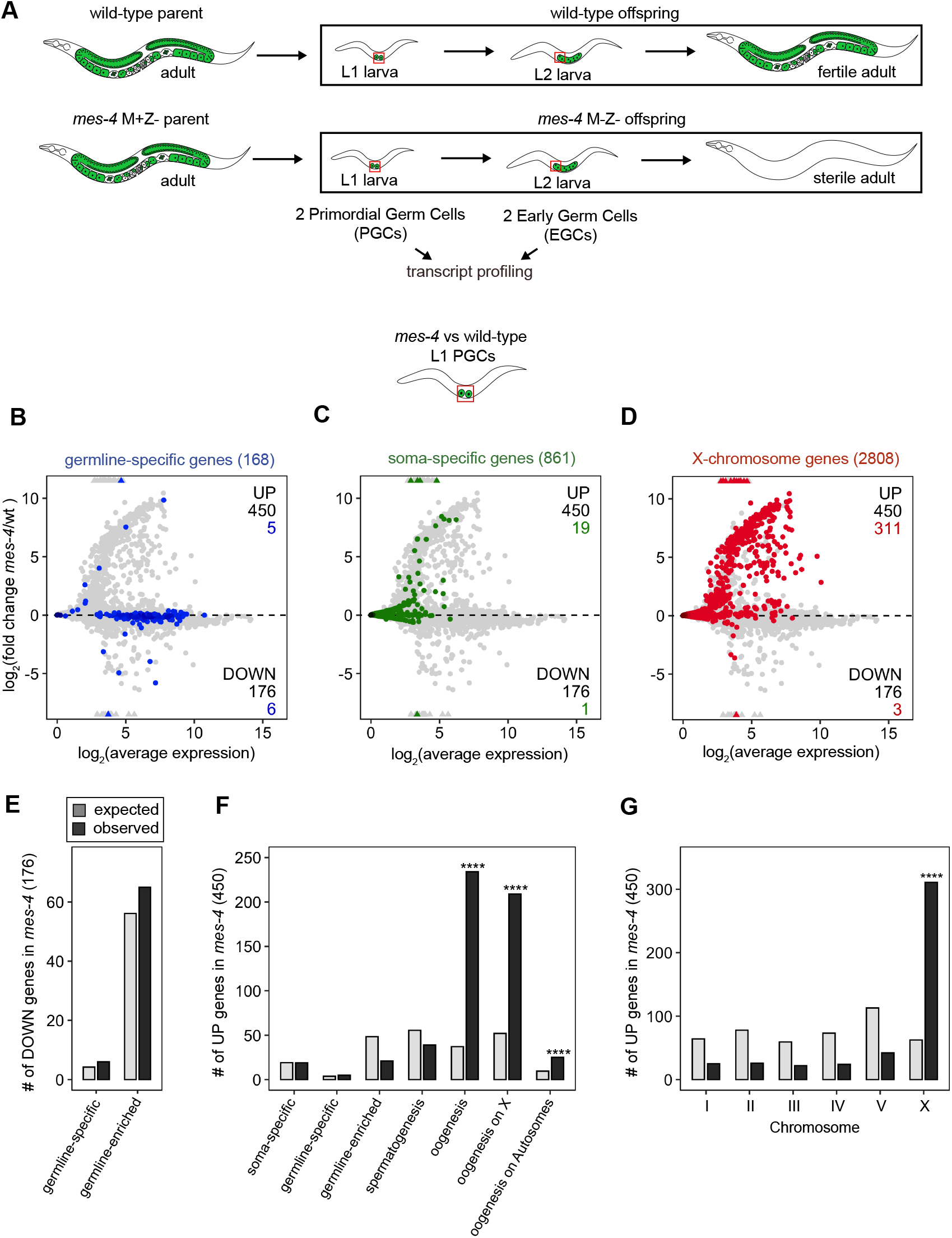
Transcriptome analysis of *mes-4* M-Z- mutant PGCs. (A) Cartoon illustrating the Maternal-effect sterile (Mes) phenotype in *mes-4* mutants. *mes-4* M+Z- (M for Maternal, Z for Zygotic) mutants cannot synthesize MES-4 (Z-) but are fertile because they inherited maternal MES-4 gene product (M+). Removal of maternal MES-4 renders *mes-4* M-Z- mutant adults sterile. Germline is green, and soma is white. Transcripts were profiled from single sets of 2 sister Primordial Germ Cells (PGCs) and from single sets of 2 Early Germ Cells (EGCs) (red boxes) hand-dissected from *mes-4* M-Z- (hereafter called *mes-4*) mutant and wild-type (wt) L1 and L2 larvae, respectively. (B,C,D) MA plots showing log_2_(average expression) versus log_2_(fold change) of transcript abundance for 20,258 protein-coding genes (circles) between *mes-4* and wt PGCs. Genes that exceed one or both plot scales (triangles) are set at the maximum value of the scale. Genes belonging to a specific gene set are colored: (B) 168 germline-specific genes are blue, (C) 861 soma-specific genes are green, and (D) 2808 X-chromosome genes are red. Differentially expressed genes in *mes-4* vs wt PGCs were identified using Wald tests in DESeq2 (Love et al., 2014) and by setting a q-value < 0.05 significance threshold. Numbers of all mis-regulated genes (black) and numbers of those in gene sets (colored) are indicated in the corners; top is upregulated (UP) and bottom is downregulated (DOWN) in *mes-4* vs wt. (E,F,G) Bar plots showing the expected number (light gray) and observed number (dark gray) of mis-regulated genes that are members of the indicated gene sets. Hypergeometric tests were performed in R to test for gene-set enrichment. P-value designations are **** < 1e-5. (E) Enrichment analyses for DOWN genes were restricted to 5,858 protein-coding genes that we defined as ‘expressed’ (minimum average read count of 1) in wt PGCs. Gene set sizes: germline-specific (140), germline-enriched (1867). (F,G) Enrichment analyses for UP genes included all 20,258 protein-coding genes in the transcriptome. Gene set sizes: soma-specific (861), germline-specific (168), germline-enriched (2176), spermatogenesis-enriched (2498), oogenesis (1671), oogenesis on X (470), oogenesis on Autosomes (1201), chrI (2888), chrII (3508), chrIII (2670), chrIV (3300), chr V (5084), chr X (2808). Figure 1—figure supplement 1: *mes-4* M-Z- EGCs have more severe transcriptome defects than *mes-4* M-Z- PGCs. Figure 1—figure supplement 2: Analysis of features of mis-regulated genes in *mes-4* M-Z- PGCs and EGCs.

To determine whether the DOWN genes include germline genes that fail to turn on normally in *mes-4* PGCs, we analyzed transcript levels and fold changes (*mes-4* vs. wt) for genes that are members of 2 ‘germline’ gene sets: 1) a ‘germline-specific’ set containing 168 genes that are expressed in germline tissue but not in somatic tissues, and 2) a ‘germline-enriched’ set containing 2176 genes that are expressed at higher levels in adults with a germline compared to adults that lack a germline (see Methods for gene sets). We found that most germline-specific genes (162 of 168 genes or 96%) and germline-enriched genes (2111 of 2176 genes or 97%) are not significantly DOWN in *mes-4* PGCs (Figure 1B). The numbers of germline-specific genes and germline-enriched genes that are DOWN are not more than expected by chance (Figure 1E).

Since some gene expression defects may not manifest until after *mes-4* PGCs have started dividing, we used our hand-dissection strategy to isolate sets of 2 PGC descendants, which we call Early Germ Cells (EGCs), from wt and *mes-4* mutant L2 larvae and profiled their transcripts. We found that *mes-4* EGCs down-regulate more germline-specific and more germline-enriched genes than *mes-4* PGCs. However, like *mes-4* PGCs, *mes-4* EGCs turn on most germline genes normally (Figure 1—figure supplement 1).

As an independent test of differential expression in PGCs, we selected 3 genes and performed smFISH to measure and compare their transcript levels between *mes-4* and wt PGCs. 2 of the genes we tested, *cpg-2* and *pgl-3,* are members of our germline-specific set and by RNA-seq analysis were DOWN or not DOWN respectively, in *mes-4* PGCs (Figure 2A-C, Figure 2—figure supplement 1). Corroborating our RNA-seq analysis, smFISH analysis showed that the average transcript abundance of *cpg-2* is significantly lower in *mes-4* vs wt PGCs, while the average transcript abundance of *pgl-3* is not significantly different (Figure 2A-C). The other gene we tested by smFISH, *chs-1*, is a member of our germline-enriched set and was consistently not mis-expressed by RNA-seq or smFISH analysis (Figure 2C, Figure 2—figure supplement 1). Together, our RNA-seq and smFISH analysis showed that *mes-4* PGCs turn on most germline genes normally.

**Figure 2:**
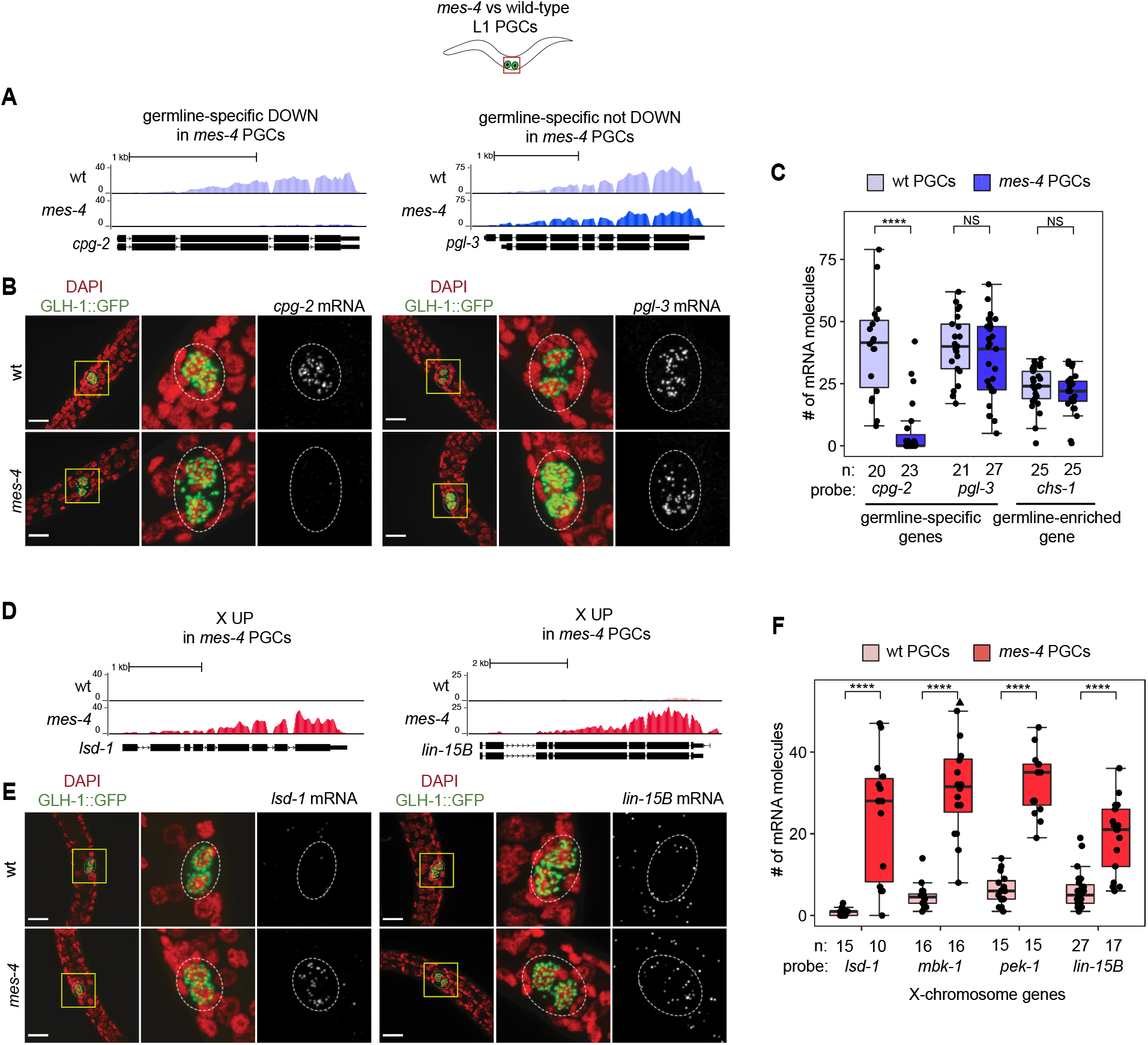
Transcript quantification in *mes-4* M-Z- mutant PGCs by single-molecule FISH (smFISH) corroborates RNA-seq results. (A,D) UCSC genome browser images showing average sequencing read coverage over Ensembl gene models for wild-type (wt) PGCs (top track) and *mes-4* PGCs (bottom track). (A) 2 germline-specific genes: *cpg-2* (left) and *pgl-3* (right). *cpg-2* is DOWN and *pgl-3* is not DOWN in transcript profiling of *mes-4* vs wt PGCs. (D) 2 X-linked UP genes in transcript profiling of *mes-4* vs wt PGCs and/or EGCs: *lsd-1* (left) and *lin-15B* (right). (B,E) Representative maximum-intensity Z-projection images of smFISH experiments in L1 larvae. DAPI-stained nuclei are red. GLH-1::GFP is green. The dashed lines circumscribe PGCs marked by GLH-1::GFP. The 2^nd^ and 3^rd^ images in each set are zoomed insets of the yellow box in the 1^st^ image. Foci in the mRNA channel (3^rd^ image in each set) represent individual transcripts. Scale bars are 10 microns. (B) smFISH RNA probes targeting *cpg-2* (left) and *pgl-3* (right) transcripts. (E) smFISH RNA probes targeting *lsd-1* (left) and *lin-15B* (right) transcripts. (C,F) Transcript quantification in smFISH 3D images of PGCs. Each circle represents 1 quantified image. The number of quantified images for each combination of probe and genotype is indicated. Boxplots show the median, the 25^th^ and 50^th^ percentiles (boxes), and the 2.5^th^ and 97.5^th^ percentiles (whiskers). Mann-Whitney tests were used to compare a gene’s transcript counts between *mes-4* and wt PGCs. P-value designations are NS > .01, **** < 1e-5. (C) Quantification of *cpg-2*, *pgl-3*, and *chs-1* transcripts. *chs-1* is in the germline-enriched gene set and is not DOWN in transcript profiling of *mes-4* vs wt PGCs. (F) Quantification of *lsd-1*, *mbk-1*, *pek-1*, and *lin-15B* transcripts, 4 X-linked UP genes in transcript profiling of *mes-4* vs wt PGCs and/or EGCs. Figure 2—figure supplement 1: Comparison of smFISH and transcript profiling data.

### MES-4 is not required to keep somatic genes off in PGCs

Chromatin regulators can protect tissue-appropriate transcription patterns by serving as a barrier to promiscuous transcription factor activity. Loss of MES-4 and PRC2 have both been shown to allow mis-expression of neuronal target genes upon ectopic expression of the transcription factor CHE-1 in the germline (Patel et al., 2012; Seelk et al., 2016). Moreover, loss of PRC2 activity in the *C. elegans* germline was recently linked to mis-expression of some neuronal genes and conversion to neuronal fate (Kaneshiro et al., 2019). Maternal MES-4 may promote offspring germline development by preventing germ cells from turning on a somatic gene expression program. To test this possibility, we examined whether UP genes in *mes-4* vs wt PGCs are members of a ‘soma-specific’ gene set that defines 861 genes expressed in soma but not in germline. We found that only 19 UP genes are soma-specific, which is not a higher number than expected by chance (Figure 1C,F). Therefore, *mes-4* PGCs do not mis-express a soma-specific program.

### MES-4 represses genes on the X chromosome including many oogenesis genes in PGCs

Repression of the X chromosomes in the *C. elegans* germline is essential for germline health (reviewed in Strome et al., 2014). We found that 311 of the 2808 (11%) protein-coding X genes are UP in *mes-4* vs wt PGCs. Strikingly, more than half of all UP genes are on the X chromosome (311 out of 450 genes), and this number is significantly higher than expected by chance (Figure 1F). We found that *mes-4* EGCs mis-express 564 X genes, including almost all of the X genes that are mis-expressed in *mes-4* PGCs and an additional 273 X genes (Figure 1—figure supplement 1C,F). These data show that *mes-4* PGCs mis-express many genes on the X chromosome and that X mis-expression becomes more severe in their descendant EGCs.

As an independent test of differential expression, we selected 4 X-linked UP genes and performed smFISH to compare their transcript levels between *mes-4* vs wt PGCs (Figure 2— figure supplement 1). smFISH analysis showed that all 4 X genes have higher transcript abundance in *mes-4* PGCs than in wt PGCs (Figure 2D-F), corroborating our transcriptome analysis. These data reveal that the X chromosome is the primary focus of MES-4 regulation in PGCs.

While most X-linked genes are repressed during germline proliferation and spermatogenesis, some are normally turned-on during oogenesis (Kelly et al., 2002; Arico et al., 2011; Tzur et al., 2018, Figure 1—figure supplement 2E). We examined whether X-linked UP genes are those that are normally turned-on during oogenesis by comparing our set of X-linked UP genes to a set of ‘oogenesis’ genes, defined as 470 X-linked and 1201 autosomal genes that are expressed at higher levels in dissected adult oogenic germlines than in dissected spermatogenic germlines (Ortiz et al., 2014). We found that 209 of the 311 (67%) X-linked UP genes and 25 of the 149 (17%) autosomal UP genes are in the oogenesis set, which are both higher numbers than expected by chance (Figure 1F). The enrichment for oogenesis X-linked genes was especially high. No other germline gene set that we tested (germline-specific, germline-enriched, and spermatogenesis) was enriched in the set of UP genes in *mes-4* PGCs (Figure 1F). Based on gene ontology (GO) analysis, the set of X-linked UP genes in *mes-4* PGCs and/or EGCs is enriched for biological process terms that characterize roles in oogenesis: ‘reproduction’ and ‘embryo development ending in birth or hatching’. (Figure 1—figure supplement 2F-H). We conclude that *mes-4* PGCs mis-express an oogenesis program involving many X-linked genes, which may interfere with the ability of mutant PGCs to proliferate.

### Mis-expression of genes on the X chromosome(s) causes germline death in *mes-4* mutants

Since X mis-expression is the largest defect to the transcriptome in *mes-4* PGCs, we hypothesized that mis-expression of the 2 X chromosomes in germlines of *mes-4* mutant hermaphrodites causes germline death. To test our hypothesis, we asked whether *mes-4* mutant males, which inherit only a single X chromosome (typically from the oocyte), have healthy germlines. We live imaged wild-type and *mes-4* mutant M-Z-hermaphrodites and males that express a germline-specific GFP reporter (GLH-1::GFP) and scored their germline health qualitatively as either ‘absent/tiny’ germline, ‘partial’ germline, or ‘full’ germline. All live-imaged *mes-4* mutant hermaphrodites lacked a germline (Figure 3A). In contrast, some *mes-4* mutant males that inherited their single X from an oocyte (X^oo^ males) had either partial or full germlines (21% and 4%, respectively). Since X chromosomes turn on during oogenesis (Kelly et al., 2002; Arico et al., 2011; Tzur et al., 2018, Figure 1—figure supplement 2E), X^oo^ males inherited an X with a history of expression. Using a *him-8* mutant, we generated wild-type and *mes-4* mutant males that instead inherited their X from a sperm (X^sp^ males), which has a history of repression because the X was not turned on previously during spermatogenesis. We tested whether *him-8; mes-4* mutant X^sp^ males that inherited a single X with a history of repression have healthier germlines than *mes-4* mutant X^oo^ males that inherited a single X with a history of expression. Strikingly, 67% *him-8; mes-4* mutant X^sp^ males made full germlines, compared to only 4% of *mes-4* mutant X^oo^ males.

**Figure 3:**
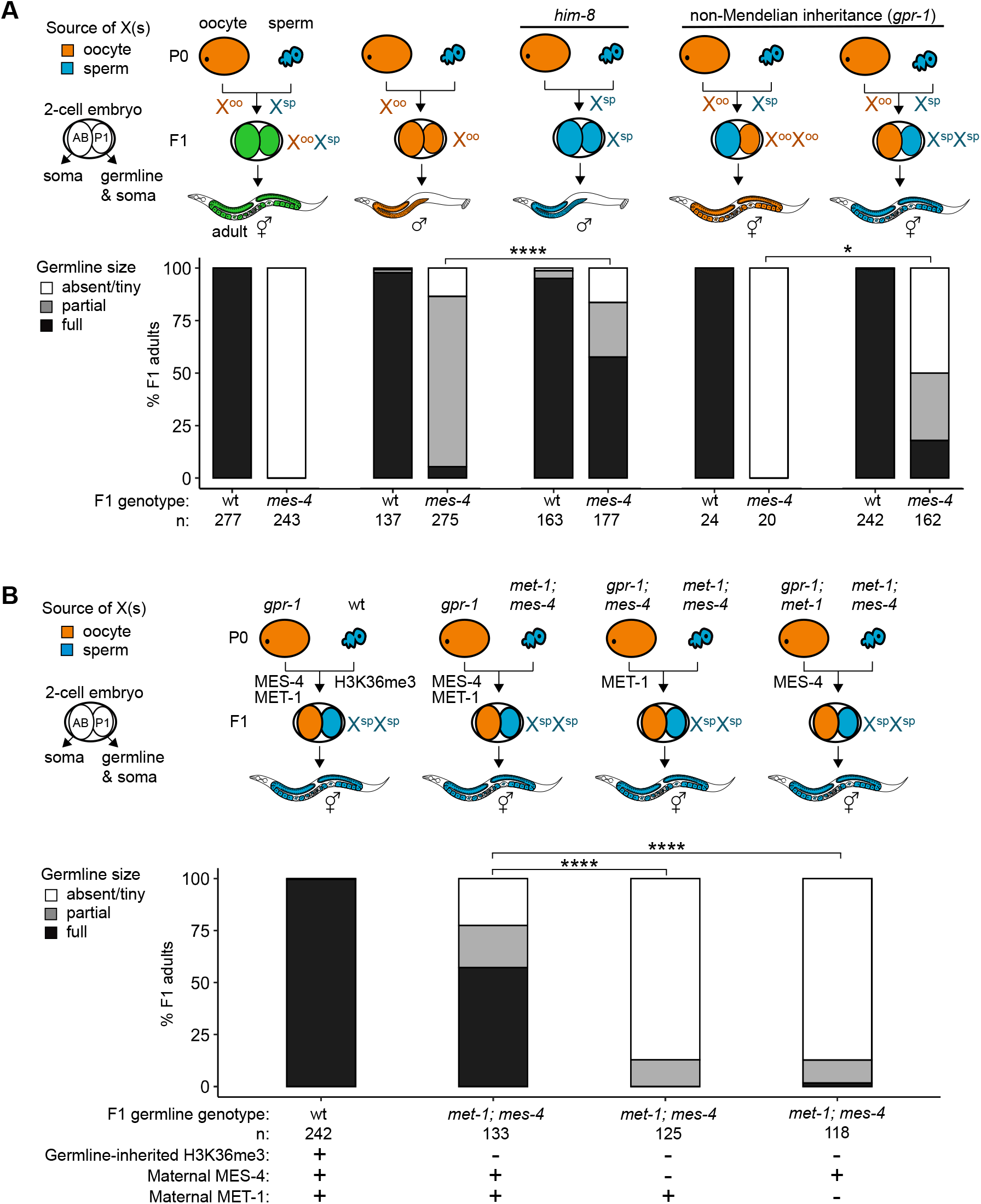
Maternally loaded MES-4 promotes germline development by repressing the X chromosomes independently from transmitting H3K36me3 across generations. (A,B) Bar plots showing distributions of germline size (absent/tiny, partial, and full) in worms with different X- chromosome compositions in their germline. Numbers of scored F1 offspring and the genotype of their germlines are indicated (*mes-4* indicates *mes-4* M-Z-). 2-cell embryos contain AB (left) and P1 (right) blastomeres. AB generates some somatic tissues; P1 generates the germline and some somatic tissues. Orange, blue, and green coloring indicate X chromosome compositions: orange is only oocyte-inherited X(s), blue is only sperm-inherited X(s), and green is 1 oocyte-inherited X and 1 sperm-inherited X. To generate ‘non-Mendelian’ F1 hermaphrodite offspring that inherited 2 genomes and therefore 2 Xs from one gamete, we mated fathers with mothers that carry a mutation in *gpr-1* (Besseling and Bringmann, 2016; Artiles et al., 2019). 2-tailed Fisher’s exact tests were used to test whether the proportion of F1 adults with a full-sized germline significantly differs between samples. P-value designations are * < 0.01 and **** < 1e-5. (A) To generate F1 male offspring that inherited their single X from the sperm, we mated parents that carry the *him-8(e1489)* allele, which causes X-chromosome nondisjunction during oogenesis in the hermaphrodite. (B) Presence or absence of sperm-inherited H3K36me3 marking, maternal MES-4, and maternal MET-1 in germlines of F1 offspring are indicated in the schematic of each cross and below each bar. Figure 3—figure supplement 1: Genetic strategies to generate and identify F1 offspring that inherited different X-chromosome endowments from parents. Figure 3—figure supplement 2: *mes-4* M-Z+ X^sp^/X^sp^ mutants do not have healthier germlines than *mes-4* M-Z- X^sp^/X^sp^ mutants. Figure 3—figure supplement 3: Further fertility analyses of *mes-4* M-Z- mutants that inherited their X(s) from the sperm.

A new and powerful genetic tool uses *gpr-1* over-expression to generate hermaphrodite worms that form a germline entirely composed of 2 genomes inherited from the sperm, or rarely 2 genomes inherited from the oocyte (Besseling and Bringmann, 2016; Artiles et al., 2019). Using this tool, we tested whether *mes-4* mutant hermaphrodites whose germline inherited from a sperm 2 X chromosomes with a history of repression (which we called X^sp^/X^sp^ hermaphrodites) have healthier germlines than *mes-4* mutant hermaphrodites whose germline inherited from an oocyte 2 X chromosomes with a history of expression (called X^oo^/X^oo^ hermaphrodites). While all *mes-4* mutant X^oo^/X^oo^ hermaphrodites lacked a germline, some *mes-4* mutant X^sp^/X^sp^ hermaphrodites had partial or full germlines (32% and 18%, respectively) (Figure 3A). Our combined genetic analysis demonstrates that mis-expression of the X chromosome(s) causes germline death in *mes-4* mutants. It also underscores that *mes-4* mutant PGCs can launch a normal germline program.

### MES-4 promotes germline health independently from its role in transmitting H3K36me3 patterns across generations

Transmission of epigenetic information across generations can impact the health of offspring. We hypothesized that maternally loaded MES-4’s role in transmitting H3K36me3 patterns from parents to offspring is essential for offspring germline development. If so, then transmission of parental chromosomes lacking H3K36me3 to offspring should cause their germline to die even if they received maternal MES-4. To test our hypothesis, we used the *gpr-1* genetic tool and the GLH-1::GFP germline marker to generate F1 adult offspring whose PGCs inherited 2 H3K36me3(-) genomes from the sperm and either did or did not inherit maternal MES-4.

Importantly, the germline in both types of F1 offspring had the same genotype (*met-1; mes-4*); the only difference between the F1s was the presence or absence of maternally loaded MES-4. We found that over half (57%) of F1 adult offspring had a full germline if their PGCs inherited 2 H3K36me3(-) genomes and maternal MES-4 (Figure 3B). In contrast, 0% of F1 adult offspring had a full germline if their PGCs inherited 2 H3K36me3(-) genomes from the sperm and did not inherit maternal MES-4 (Figure 3B). This result shows that maternally loaded MES-4 is critical for offspring germline development but that its critical role is not to transmit H3K36me3 patterns from parents to offspring.

Presence of maternal MES-4 allows many F1 offspring whose PGCs inherited 2 H3K36me3(-) genomes from the sperm to make a full germline. One possibility is that maternally loaded H3K36 HMTs can re-establish sufficient levels of H3K36me3 marking to H3K36me3(-) chromosomes in PGCs for germline development. Since MES-4 cannot catalyze de novo H3K36me3 marking on H3K36me3(-) chromosomes (Furuhashi et al., 2010; Kreher et al., 2018), re-establishment of H3K36me3 levels would require the other H3K36 HMT MET-1, which can catalyze de novo marking in response to transcriptional turn-on, like H3K36 HMTs in other species. We found that removal of maternal MET-1, like removal of maternal MES-4, caused almost all F1 offspring whose PGCs inherited 2 H3K36me3(-) genomes from sperm to lack a germline. These findings show that maternal loads of both H3K36 HMTs are required for F1 offspring whose PGCs inherited 2 H3K36me3(-) genomes from sperm to make a germline, and suggest that newly established H3K36me3 marking of H3K36me3(-) chromosomes by maternally loaded HMTs can enable PGCs to make a germline.

### LIN-15B causes X mis-expression in germlines of *mes-4* M+Z- adults

MES-4 levels are low on the X chromosome(s) (Bender et al., 2006; Rechtsteiner et al., 2010; Furuhashi et al., 2010; Gaydos et al., 2012; Kreher et al., 2018). We therefore hypothesized that MES-4 represses X genes indirectly by regulating the expression or activity of 1 or more downstream factor(s). For example, MES-4’s activity on autosomes may repress X genes by concentrating a transcriptional repressor onto the X or by sequestering a transcriptional activator away from the X (Gaydos et al., 2012; Cabianca et al., 2019; Georgescu et al., 2020). Several lines of evidence made the THAP transcription factor LIN-15B a strong candidate for causing X mis-expression in germlines that lack MES-4. First, our analysis of publicly available LIN-15B ChIP data from whole embryos and larvae found that LIN-15B targets the promoter of many X genes that are repressed by MES-4 in PGCs and/or EGCs (Figure 4 -- Supplement 1). Second, *lin-15B* is X-linked and UP in *mes-4* vs wt PGCs (Figure 2E,F and Figure 2—figure supplement 1). Third, LIN-15B has been reported to promote expression of X-linked genes in PGCs and adult germlines (Lee et al., 2017; Robert et al., 2020). Finally, *mes-4* and *lin-15B* genetically interact in somatic cells to control expression of germline genes such as genes encoding P-granule components (Petrella et al., 2011).

We hypothesized that LIN-15B causes mis-expression of X genes in germlines that lack MES-4. To test our hypothesis, we used RNA-seq to determine whether gonads dissected from *mes-4* M+Z-*; lin-15B* M-Z- double mutant adults have reduced levels of X mis-expression compared to gonads dissected from *mes-4* M+Z- single mutant adults. Our differential expression analyses showed that *mes-4* M+Z-*; lin-15B* M-Z- adult gonads up-regulate considerably fewer X genes (112 X genes) than *mes-4* M+Z- adult gonads (367 X genes) (Figure 4A and Figure 4—figure supplement 2A). Furthermore, the 367 X-UP genes in *mes-4* M+Z- adult gonads had closer-to-wild-type transcript levels in *mes-4* M+Z-*; lin-15B* M-Z- adult gonads (Figure 4B), and 323 of those X-UP genes were scored as X-DOWN in *mes-4* M+Z-*; lin-15B* M-Z- compared to *mes-4* M+Z- (Figure 4C and Figure 4—figure supplement 1C). Our results show that LIN-15B is responsible for much of the X mis-expression in *mes-4* M+Z- adult gonads.

**Figure 4:**
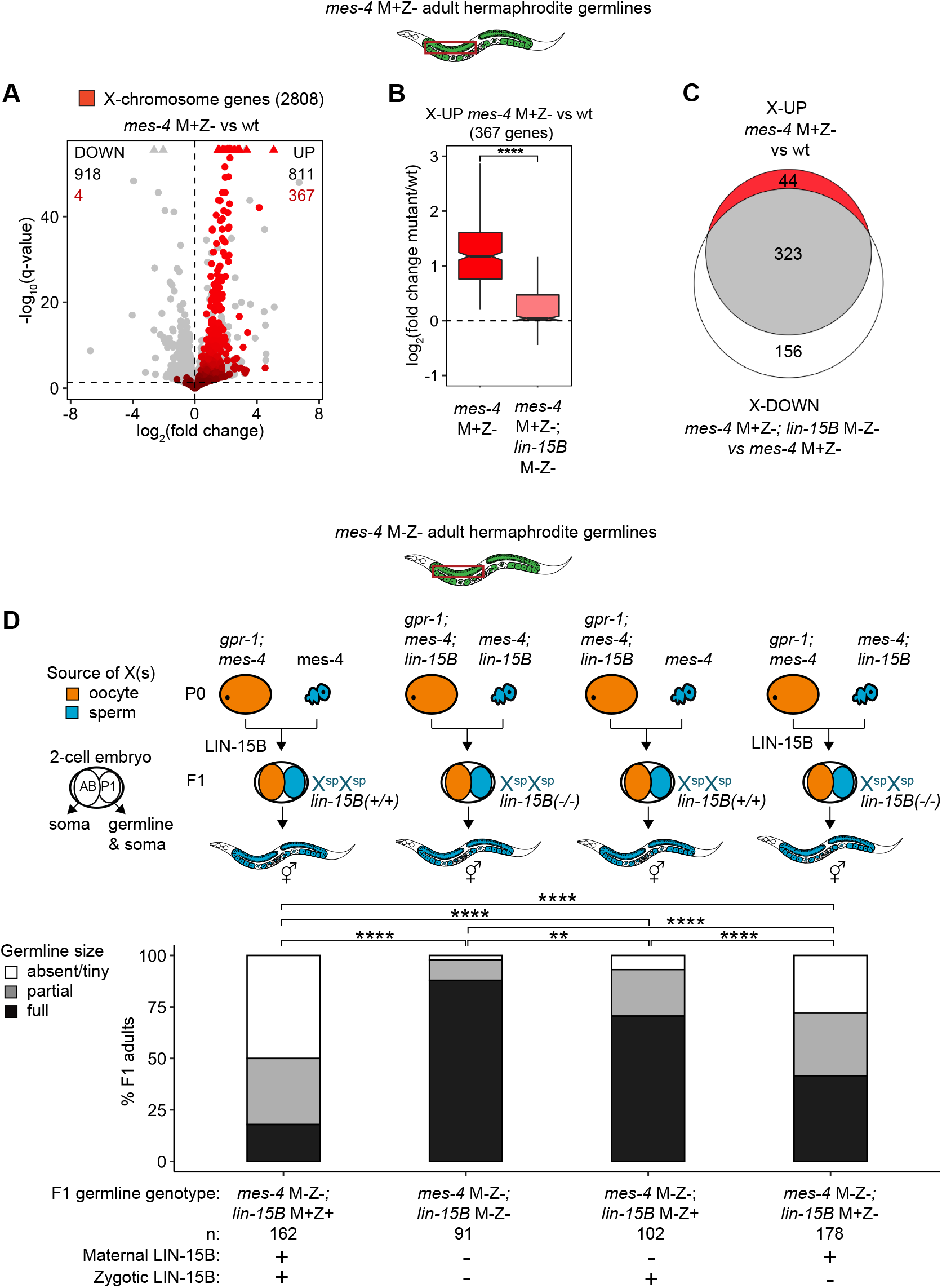
Loss of LIN-15B reduces X mis-expression in *mes-4* M+Z- adult germlines and suppresses germline death in *mes-4 M-Z-* mutants. (A) Volcano plot showing log_2_(fold change) of transcript abundance and significance [log_10_(q-value)] for 20,258 protein-coding genes (circles) between gonads dissected from *mes-4* M+Z- vs wild-type (wt) adults. Genes that exceed the plot scale (triangles) are set at the maximum value of the scale. X-chromosome genes (2808) are red. Genes above the horizontal line (q-value of 0.05) are considered significantly mis-regulated. The number of all mis-regulated protein-coding genes (black) and the number of those that are X-linked (red) are indicated in the corners; left is downregulated (DOWN) and right is upregulated (UP) in *mes-4* M+Z*-* vs wt. (B) Boxplots showing log_2_(fold change) in transcript abundance for the 367 X-UP genes in *mes-4* M+Z- vs wt gonads between *mes-4* M+Z- vs wt gonads (red) and between *mes-4* M+Z-*; lin-15B* M-Z- vs wt gonads (pink). Boxplots show the median, the 25^th^ and 50^th^ percentiles (boxes), and the 2.5^th^ and 97.5^th^ percentiles (whiskers). Waists around the median indicate 95% confidence intervals. Mann- Whitney tests were used to compare samples. (C) Venn diagram comparing the 367 X-UP genes in *mes-4* M+Z- vs wt and the 479 X-DOWN genes in *mes-4* M+Z-*; lin-15B* M-Z- vs *mes-4* M+Z- gonads. (D) Bar plots as described in the legend of Figure 3. Genotypes of hermaphrodite and male parents are indicated at the top. All scored F1 offspring are non-Mendelian segregants (caused by the *gpr-1* mutation in mother worms) whose germline inherited 2 genomes and therefore 2 Xs from the sperm. The F1 germline’s genotype with respect to *lin-15B* is indicated to the right of the 2-cell embryos; ‘+’ is wild-type allele, ‘-‘ is null allele. The presence or absence of maternal LIN-15B and zygotic LIN-15B in the germline of F1 offspring is indicated in the schematic of each cross and below each bar. 2-sided Fisher’s exact tests were used to test whether the proportion of F1 adults with a full-sized germline significantly differed between samples. P-value designations are ** < 0.001, **** < 1e-5. Figure 4—figure supplement 1: Identification and testing of candidate transcription factors for a role in causing sterility of *mes-4* M-Z- mutants. Figure 4—figure supplement 2: Further analysis of how LIN-15B impacts the transcriptome of *mes-4* M+Z- dissected adult germlines. Figure 4—figure supplement 3: Removal of LIN-15B improves germline health in *mes-4* M-Z- X^oo^/X^sp^ mutant hermaphrodites.

### LIN-15B causes sterility in *mes-4* M-Z- mutants

Since LIN-15B causes X mis-expression in *mes-4* M+Z- adult gonads, we hypothesized that removal of LIN-15B would allow *mes-4* M-Z- mutants to make healthier germlines. We compared germline health in *mes-4* M-Z- mutant hermaphrodite adults with an X^sp^/X^sp^ germline, comparing those that had maternal and zygotic LIN-15B (*lin-15B* M+Z+) and those X^sp^/X^sp^ mutant hermaphrodite adults and those that lacked either maternal LIN-15B (*lin-15B* M- Z+), zygotically synthesized LIN-15B (*lin-15B* M+Z-), or both (*lin-15B* M-Z-). We found that removal of either maternal LIN-15B, zygotic LIN-15B, or both caused more *mes-4* M-Z- X^sp^/X^sp^ germlines to be full-sized (Figure 4D). Notably, loss of maternal LIN-15B caused better recovery of germline health than loss of zygotic LIN-15B, and loss of both had an additive effect. Strikingly, 88% of *mes-4* M-Z-; *lin-15B* M-Z- X^sp^/X^sp^ germlines were full-sized. We conclude that both maternal and zygotic sources of the transcription factor LIN-15B cause germline loss in *mes-4* M-Z- mutants.

Removal of LIN-15B may only allow *mes-4* M-Z- mutant hermaphrodites to make a full-sized germline if that germline inherited 2 X chromosomes with a history of repression (from sperm), which by itself improves germline health in *mes-4* M-Z- mutants (Figure 3A, Figure 4D). We analyzed the impact of loss of LIN-15B on germline health in X^oo^/X^sp^ mutants that inherited 1 of their 2 X chromosomes with a history of expression (from the oocyte). We found that 29% of *mes-4* M-Z-; *lin-15B* M-Z- X^oo^/X^sp^ adult mutant hermaphrodites made full-sized germlines compared to 0% of *mes-4* M-Z-; *lin-15B* M+Z+ adult mutant hermaphrodites (Figure 4—figure supplement 3). This finding demonstrates that removal of LIN-15B can even allow *mes-4* M-Z- X^oo^/X^sp^ mutants to make a full-sized germline.

To investigate whether other factors contribute to sterility in *mes-4* M-Z- mutants, we identified candidate genes that met 1 or more of 3 criteria: 1) they are X-linked and UP in *mes-4* PGCs and/or EGCs, 2) there is evidence of them binding to the promoter region of at least 25% of X-linked UP genes, and 3) they target a DNA motif that is enriched in the promoter of X-linked UP genes (Figure 4—figure supplement 1). We identified 20 top candidates based on the above criteria plus 4 histone acetyltransferases (HATs) that are involved in transcriptional activation and tested whether their depletion by RNAi causes *mes-4* M-Z- mutants to make healthier germlines. Of the 21 genes tested, only RNAi depletion of LIN-15B caused *mes-4* M-Z- mutants to make healthier germlines (Figure 4—figure supplement 1). We conclude that LIN- 15B is a major contributor to germline death in *mes-4* M-Z- mutants.

### MES-4 cooperates with the chromatin regulator PRC2 to repress X genes

In addition to MES-4, the maternally loaded H3K27 HMT Polycomb Repressive Complex 2 (PRC2), composed of MES-2, MES-3, and MES-6, promotes germline survival and development by repressing genes on the X chromosome (Gaydos et al., 2012; Gaydos et al., 2014). To test if MES-4 and PRC2 cooperate to protect germline health by repressing a similar set of X genes, we compared transcript profiles in wild-type (wt), *mes-3* M-Z-, and *mes-4* M-Z- PGCs and EGCs. In Principal Component Analysis (PCA), the top 2 principal components captured 41% of the variance across all samples and clustered *mes-4* and *mes-3* mutant samples together by germline stage and away from wild-type samples (Figure 5A). Using differential expression analysis, we identified 354 X genes UP in *mes-3* vs wt PGCs and 443 X genes UP in *mes-3* vs wt EGCs. We found that stage-matched *mes-3* and *mes-4* samples up-regulate a highly similar set of X genes (Figure 5B-C). Next, we compared log_2_(fold change) (mutant vs wt) of mis-regulated X genes between *mes-4* and *mes-3* PGCs and between *mes-4* and *mes-3* EGCs. We found a positive, albeit small, correlation between PGCs (0.22 Spearman’s correlation coefficient) and a stronger correlation between EGCs (0.44 Spearman’s correlation coefficient) (Figure 5D-E). Moreover, as in *mes-4* M-Z- mutants, loss of LIN-15B caused *mes-3* M-Z- mutants to make healthier germlines (Figure 4—figure supplement 3). We conclude that MES-4 and PRC2 cooperate to ensure germline survival in M-Z- mutant larvae by repressing similar sets of X genes and that both operate through LIN-15B.

**Figure 5:**
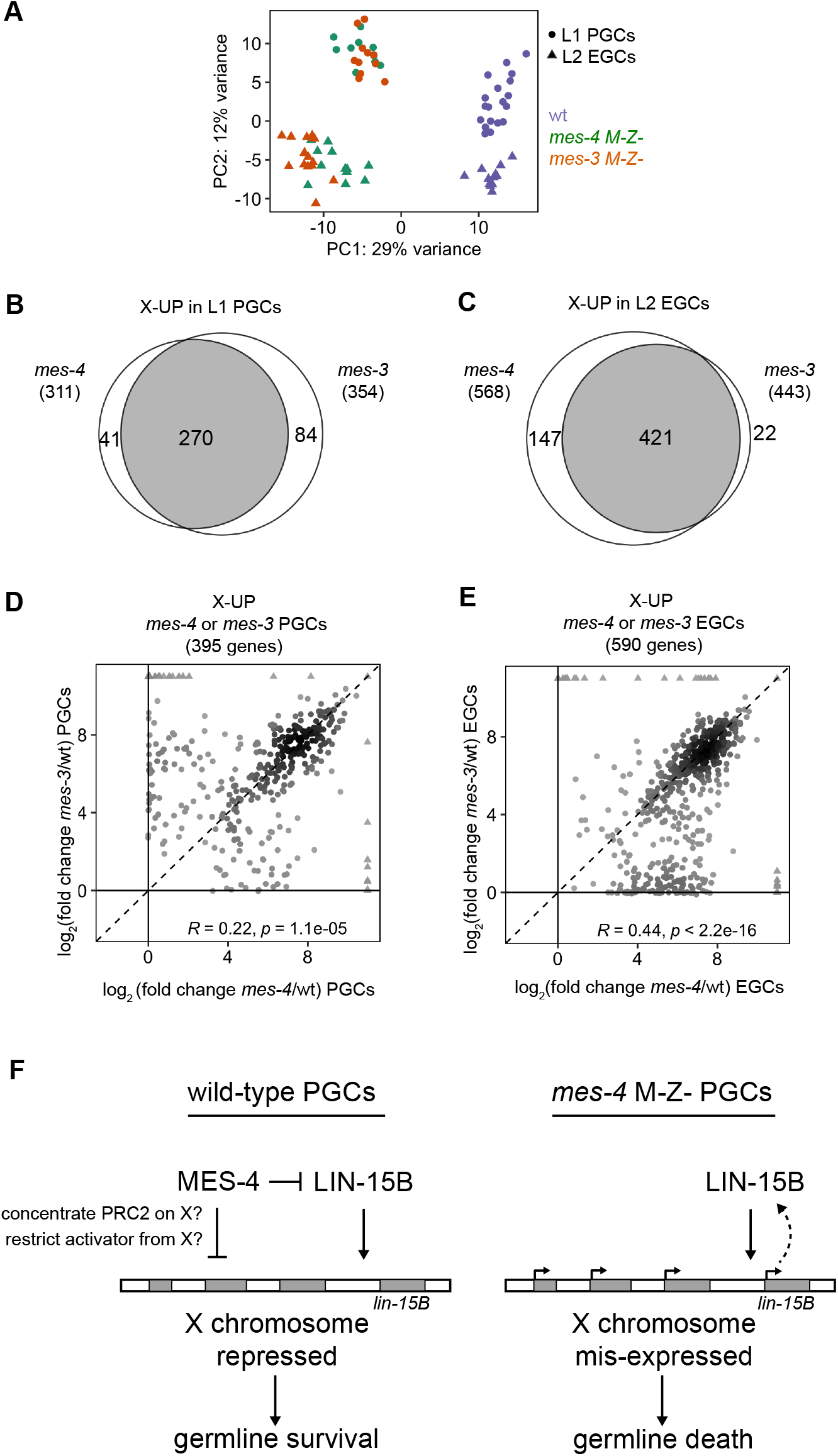
*mes-4* M-Z- and *mes-3* M-Z- nascent germlines mis-express a highly similar set of X genes. (A) Principal Component Analysis (PCA) including all replicates of wt (blue), *mes-4* M-Z- (green), and *mes-3* M-Z- (dark orange) PGCs (circles) and EGCs (triangles). The percentages of total variance across all samples described by the top 2 principal components are indicated. (B,C) Venn diagrams comparing X-UP genes (mutant vs wt) in *mes-4* and *mes-3* PGCs (B) and in *mes-4* and *mes-3* EGCs (C). (D,E) Scatterplots comparing log_2_(fold change) (mutant vs wt) of transcript abundance for X-UP genes (circles) in *mes-4* or *mes-3* PGCs (D) or in *mes-4* or *mes-3* EGCs (E). The Spearman correlation coefficient along with its p-value is indicated at the top of each scatterplot. (F) Cartoon model illustrating how MES-4 protects germline survival by repressing X genes (gray boxes). MES-4 may indirectly repress X genes, including *lin-15B*, by concentrating a repressor (e.g. PRC2) on the X or by restricting an activator (e.g. histone acetyltransferase or LIN-15B) from the X. Our findings identify LIN-15B as a key player in activating X genes and causing germline death upon loss of MES-4. LIN-15B may activate X genes directly by binding to those genes or indirectly by regulating 1 or more other transcription factors. Figure 5—figure supplement 1: Comparison of X mis-expression in PGCs and EGCs dissected from various chromatin regulator mutants.

The chromodomain protein MRG-1 is a candidate reader and effector of H3K36me3 that, like MES-4 and PRC2, promotes germline development by repressing X genes (Fujita et al., 2002; Takasaki et al., 2007). To test if MRG-1 represses the same set of X genes as MES-4 and PRC2, we profiled transcripts from PGCs hand-dissected from *mrg-1* M-Z- L1 larvae. We found that *mrg-1* PGCs mis-express 440 X genes, 225 of which are also mis-expressed in *mes-4* and *mes-3* PGCs and EGCs (Figure 5—figure supplement 1). These findings add MRG-1 to the team of maternal regulators that ensure PGC survival and health by repressing the X.

## DISCUSSION

In this study, we investigated how a maternally supplied chromatin regulator protects germline immortality and promotes germline health. We found that nascent *C. elegans* germlines (PGCs and EGCs) that completely lack maternal MES-4 mis-express over a thousand genes, most of which are on the X chromosome. We further demonstrated that X mis-expression is the cause of germline death in *mes-4* M-Z- mutants. Removal of a single transcription factor, LIN-15B, reduced X mis-expression in the germline of *mes-4* mutant mothers (*mes-4* M+Z-) and was sufficient to allow most of their offspring (*mes-4* M-Z-) to develop full-sized germlines. Intriguingly, *lin-15B* is itself X-linked and mis-expressed in nascent germlines that lack MES-4, highlighting *lin-15B* as a key target for MES-4 repression. We favor a model where maternal MES-4 promotes offspring germline development by preventing LIN-15B from activating a germline-toxic program of gene expression from the X chromosome (Figure 5F). This work underscores how maternally supplied factors can guide development of specific tissues in offspring by protecting their transcriptome.

Maternal MES-4 binds to ∼5000 genes in embryos (Rechtsteiner et al., 2010), many of which were previously expressed in the maternal germline and need to be expressed in the offspring germline. Yet surprisingly, lack of MES-4 does not impact the expression of most germline genes in PGCs. Furthermore, our genetic findings show that *mes-4* mutants can develop a full-sized germline if they inherit X chromosomes that have a history of repression, demonstrating that MES-4 is not required for PGCs to launch a germline program. Interestingly, MES-4 is required for the mis-expression of germline genes in somatic tissues of several mutants, such as *spr-5; met-2*, *lin-15B*, and mutants of DREAM complex components (Wang et al., 2005; Petrella et al., 2011; Carpenter et al., 2021). This suggests that maternal MES-4 has tissue-dependent roles in gene regulation. Such context-dependent roles may be mediated by different H3K36me3 ‘reader’ complexes, as observed in other organisms (Yochum and Ayer, 2002; Cai et al., 2003; Chen et al., 2009). Our findings clarify the role of maternal MES-4 in regulating the transcriptome of newborn germlines.

There has been a concerted effort in recent years to determine how epigenetic inheritance impacts offspring health (e.g. Heard and Martienssen, 2014; Klosin et al., 2017; Tabuchi et al., 2018; Kaneshiro et al., 2019; Fitz-James and Cavalli, 2022). Maternal MES-4’s role in propagating gamete-inherited H3K36me3 patterns through embryogenesis is a clear example of epigenetic inheritance (Kreher et al., 2018). By taking advantage of the *gpr-1* genetic tool, we demonstrated that inheritance of H3K36me3 patterns from parents is not required for offspring germline development. However, additional loss of either maternal MES-4 or maternal MET-1 (the two H3K36 HMTs in *C. elegans*) rendered almost all worms sterile, suggesting an important role for H3K36me3 marking during early germline development. We speculate that maternal MES-4 and maternal MET-1 cooperate to allow germline development by restoring sufficient levels and proper patterns of H3K36me3 marking to chromosomes inherited lacking H3K36me3. In this scenario, we envision that maternal MET-1 first catalyzes new H3K36me3 marking on genes co-transcriptionally during the first wave of zygotic genome activation in PGCs (Furuhashi et al., 2010; Kreher et al., 2018), after which maternal MES-4 maintains MET-1-generated patterns of H3K36me3 through early germline development to prevent germline death. The importance of H3K36me3 marking is also highlighted by our finding that loss of the candidate H3K36me3 ‘reader’ MRG-1 (homolog of yeast Eaf-3, fly MSL3, and human MRG15) (Gorman et al., 1995; Eisen et al., 2001; Cai et al., 2003; Bertram and Pereira-Smith, 2001; Joshi and Struhl, 2005) causes PGCs to mis-express a set of X genes similar to that caused by loss MES-4 and also causes death of the nascent germline.

Since MES-4 binding and its HMT activity are very low across almost the entire X chromosome (Rechtsteiner et al., 2010), it is likely that MES-4 regulates expression of X genes indirectly in PGCs. One possible mechanism for indirect regulation is that MES-4 generates H3K36me3 on autosomes to repel and concentrate a transcriptional repressor on the X chromosome(s). An attractive candidate repressor is the H3K27 HMT Polycomb Repressive Complex 2 (PRC2): in germlines, PRC2 activity is concentrated on the X chromosome(s) (Bender et al., 2004), PRC2 represses a highly similar set of X genes as MES-4 (this work; Gaydos et al., 2012; Lee et al., 2017), and loss of PRC2 causes maternal-effect sterility, like loss of MES-4 (Capowski et al., 1991). H3K36me3’s role in antagonizing methylation of H3K27 is well documented (Schmitges et al., 2011; Yuan et al., 2011; Gaydos et al., 2012; Evans et al., 2016). An alternative possibility is that H3K36me3 in germlines sequesters a transcriptional activator on autosomes and away from the X chromosome(s) as it does to the histone acetyltransferase (HAT) CBP-1 in *C. elegans* intestinal cells (Cabianca et al., 2019; Georgescu et al., 2020).

We identified the THAP transcription factor LIN-15B as a cause of X mis-expression in fertile germlines that lack MES-4 (*mes-4* M+Z- mutant mothers) and a major driver of germline death in their *mes-4* M-Z- mutant offspring. Interestingly, *lin-15B* is itself an X-linked gene that is mis- expressed in *mes-4* M-Z- mutant PGCs and EGCs. This suggests that although maternal MES-4 regulates expression of thousands of genes in offspring germlines, it only needs to repress *lin- 15B* to allow germline survival. However, there are likely additional factors besides LIN-15B that contribute to X mis-expression and germline death in *mes-4* M-Z- mutants, as removal of LIN- 15B does not allow all *mes-4* M-Z- mutants to make full germlines. We tested whether RNAi depletion of 21 candidate transcription factors and HATs improves germline health in *mes-4* mutants; we found no hits other than LIN-15B. Recent studies found that upregulation of *lin-15B* also causes sterility in *nanos* mutants and *set-2* (H3K4 HMT) mutants and leads to up-regulation of other X genes (Lee et al., 2017; Robert et al., 2020). Those findings coupled with ours suggest that excessive LIN-15B activity causes germline-toxic levels of X-chromosome expression and that the germline uses multiple protective mechanisms to antagonize LIN-15B.

Why is X mis-expression toxic to the germline? Our transcriptome analysis found that the X genes mis-expressed in *mes-4* M-Z- nascent germlines are highly enriched for genes that are normally turned on during oogenesis. This suggests the intriguing possibility that maternal MES- 4 promotes germline development of PGCs by antagonizing an oogenesis program that interferes with the proliferative fate of PGCs. Since the oocyte-inherited X chromosome has a history of expression, it may be prone to turning on in, and thereby causing death of, offspring PGCs that lack MES-4. In support of this, offspring PGCs that lack MES-4 can develop into full-sized germlines if they inherit only X chromosomes that have a history of repression (from the sperm). Our findings support a model where MES-4 prevents activation of the oocyte-inherited X chromosome in PGCs by opposing transcription factors such as LIN-15B.

How LIN-15B causes X mis-expression in germlines that lack MES-4 is unclear. Several studies have focused on LIN-15B’s role as a repressor of germline genes in somatic tissues (Wang et al., 2005; Petrella et al., 2011; Wu et al., 2012). Recently, LIN-15B was shown to promote repressive H3K9me2 marking in the promoter of germline-specific genes in somatic cells (Rechtsteiner et al., 2019). In germlines, LIN-15B may activate expression of X genes indirectly, e.g. by downregulating or antagonizing a repressor of X genes. Alternatively, LIN-15B may have context-dependent roles in gene expression, a well-known feature of many transcription factors (Fry and Farnham, 1999), and may directly activate expression of X genes. In support of this model, our analysis of LIN-15B ChIP data from whole embryos and larvae found that LIN-15B is associated with the promoter of many X genes that are repressed by MES-4 in newborn germlines. Clarification of LIN-15B’s mode of action in germlines requires analysis of germline-specific chromatin patterns of LIN-15B binding, biochemical experiments, and identification of LIN-15B’s functional partners.

An outstanding question is what launches the germline program in *C. elegans* PGCs (Strome and Updike, 2015). MES-4 has been a prime candidate since it transmits H3K36me marking of germline genes from parent germ cells to offspring germ cells and so has been invoked as passing a ‘memory of germline’ across generations. Our findings disprove that “memory of germline” role for MES-4, since *mes-4* mutant PGCs turn on most germline genes normally and can undergo normal germline development if they inherit X chromosomes with a history of repression. Other attractive contenders for specifying the germline fate of PGCs have been germ granules and small RNAs. Several studies demonstrated that *C. elegans* germ granules (aka P granules) protect germline fate but are not needed to specify it (Gallo et al., 2010; Updike et al., 2014; Knutson et al., 2017). Among small RNAs, 22G small RNAs (22 bp long and starting with a G) are particularly attractive as possible germline determinants. They are bound to the argonaute CSR-1 and have been shown to target most germline-expressed genes and promote expression of some (Claycomb et al., 2009; Conine et al., 2010; Wedeles et al., 2013; Cecere et al., 2014). Complete loss of CSR-1 or DRH-3, an RNA helicase that generates 22G RNAs, causes sterility (Duchaine et al., 2006; Claycomb et al., 2009; Gu et al., 2009). However, hypomorphic mutations in the helicase domain of DRH-3 that abolish production of most 22G RNAs do not impact germline formation, suggesting that 22G RNAs are not needed to specify germline fate (Gu et al., 2009). We propose the intriguing possibility that in *C. elegans*, germline fate is the default, which must be protected in the germline (e.g. by MES proteins and P granules) and opposed in somatic tissues (e.g. by DREAM and LIN-15B).

## MATERIALS AND METHODS

### Worm strains

All worms were maintained at 20°C on Nematode Growth Medium (NGM) plates spotted with *E. coli* OP50 (Brenner, 1974). Strains generated (*) and used in this study are listed below. GLH-1 (Vasa) is a component of germ granules and is specifically and highly expressed in germline cells; *glh-1::GFP* was engineered into numerous strains to mark germ cells.

**DUP64** *glh-1(sams24[glh-1::GFP::3xFLAG]) I*
**SS1491**\* *glh-1(sams24[glh-1::GFP::3xFLAG]) I; mes-4(bn73)/tmC12[egl-9(tmIs1194)] V*
**SS1293\*** *glh-1(sams24[glh-1::GFP::3xFLAG]) I; mrg-1(qa6200)/qC1[dpy-19(e1259) glp-1(q339)] III*
**SS1492**\* *mes-3(bn199)/tmC20 [unc-14(tmIs1219) dpy-5(tm9715)] glh-1(sams24[glh-1::GFP::3xFLAG]) I*
**SS1476**\* *met-1(bn200)/tmC20[unc-14(tmIs1219) dpy-5(tm9715)] glh-1(sams24[glh-1::GFP::3xFLAG]) I*
**SS1494**\* *met-1(bn200)/tmC20[unc-14(tmIs1219) dpy-5(tm9715)] glh-1(sams24[glh-1::GFP::3xFLAG]) I; mes-4(bn73)/tmC12[egl-9(tmIs1197)] V*
**SS1497**\* *glh-1(sams24[glh-1::GFP::3xFLAG]); oxTi421[eft-3p::mCherry::tbb-2 3’UTR + Cbr-unc-119(+)] X*
**SS1514**\* *glh-1(sams24[glh-1::GFP::3xFLAG]); mes-4(bn73)/tmC12[egl-9(tmIs1194)] V; oxTi421[eft-3p::mCherry::tbb-2 3’UTR + Cbr-unc-119(+)] X*
**SS1503**\* *glh-1(sams24[glh-1::GFP::3xFLAG]) I; him-8(e1489) IV*
**SS1500**\* - *glh-1(sams24[glh-1::GFP::3xFLAG]) I; him-8(e1489) IV*; *mes-4(bn73)/tmC12[egl-9(tmIs1194)] V*
**SS1498**\* *glh-1(sams24[glh-1::GFP::3xFLAG]) I; him-8(e1489) IV; oxTi421[eft-3p::mCherry::tbb-2 3’UTR + Cbr-unc-119(+)] X*
**SS1493**\* *glh-1(sams24[glh-1::GFP::3xFLAG]); him-8(e1489) IV; mes-4(bn73)/tmC12[egl-9(tmIs1194)] V; oxTi421[eft-3p::mCherry::tbb-2 3’UTR + Cbr-unc-119(+)] X*
**SS1515**\* *glh-1(sams24[glh-1::GFP::3xFLAG]) I; ccTi1594[mex-5p::GFP::gpr-1::smu-1 3’UTR + Cbr-unc-119(+), III: 680195] III; hjSi20[myo-2p::mCherry::unc-54 3’UTR] IV*
**SS1516**\* *glh-1(sams24[glh-1::GFP::3xFLAG]) I; ccTi1594[mex-5p::GFP::gpr-1::smu-1 3’UTR + Cbr-unc-119(+), III: 680195] III; hjSi20[myo-2p::mCherry::unc-54 3’UTR] IV; mes-4(bn73)/tmC12[egl- 9(tmIs1194)] V*
**SS1517**\* *glh-1(sams24[glh-1::GFP::3xFLAG]) I; ccTi1594[mex-5p::GFP::gpr-1::smu-1 3’UTR + Cbr-unc-119(+), III: 680195] III; hjSi20[myo-2p::mCherry::unc-54 3’UTR] IV; mes-4(bn73)/tmC12[egl- 9(tmIs1194)] V; lin-15B(n744) X*

**SS1518**\* *met-1(bn200) glh-1(sams24[glh-1::GFP::3xFLAG]) I; ccTi1594[mex-5p::GFP::gpr-1::smu- 1 3’UTR + Cbr-unc-119(+), III: 680195] III. hjSi20[myo-2p::mCherry::unc-54 3’UTR] IV*

**SS1511**\* *glh-1(sams24[glh-1::GFP::3xFLAG]) I; mes-4(bn73)/tmC12[egl-9(tmIs1194)] V; lin- 15B(n744) X*

**SS1512**\* *mes-3(bn199)/tmC20[unc-14(tmIs1219) dpy-5(tm9715)] glh-1(sams24[glh- 1::GFP::3xFLAG]) I; lin-15B(n744) X*

### Creation of *mes-3* and *met-1* null alleles by CRISPR-Cas9

The null alleles *mes-3(bn199)* and *met-1(bn200)* linked to *glh-1::GFP* were created by inserting TAACTAACTAAAGATCT into the 1st exon of each locus. The resulting genomic edit introduced a TAA stop codon in each reading frame, a frame shift in the coding sequence, and a BglII restriction site (AGATCT) for genotyping. Alt-R crRNA oligos (IDT) were designed using CRISPOR (Concordet and Haeussler, 2018) and the UCSC Genome Browser (ce10) to produce highly efficient and specific Cas9 cleavage in the 1st exons of *mes-3* and *met-1*. Ultramer ssDNA oligos (IDT) containing 50 bp micro-homology arms were used as repair templates. A *dpy-10* co-CRISPR strategy (Arribere et al., 2014) was used to isolate strains carrying our desired mutations. Briefly, 2.0 uL of 100 µM *mes-3* or *met-1* crRNA and 0.5 uL of 100 µM *dpy-10* cRNA were annealed to 2.5 uL of 100 µM tracrRNA (IDT) by incubation at 95°C for 2 minutes, then at room temperature for 5 minutes, to produce sgRNAs. sgRNAs were complexed with 5 uL of 40 µM Cas9 protein at room temperature for 5 minutes, 1 uL of 40 µM *mes-3* or *met-1* repair template and 1 uL of 40 µM *dpy-10(cn64)* repair template were added, and the mix was centrifuged at 13,000g for 10 minutes. All RNA oligos were resuspended in duplex buffer (IDT, #11-05-01-03). Mixes were injected into 1 or both gonad arms of ∼30 DUP64 adults. Transformant progeny were isolated and back-crossed 4x to DUP64.

### Isolation of single sets of 2 sister PGCs or 2 EGCs

L1 larvae hatched within a 30-minute window in the absence of food were allowed to feed for 30 minutes to start PGC development. Larvae were partially immobilized in 15 uL drops of egg buffer (25 mM HEPES, pH 7.5, 118 mM NaCl, 48 mM KCl, 2 mM MgCl_2_, 2 mM CaCl_2_, adjusted to 340 mOsm) on poly-lysine coated microscope slides and hand-dissected using 30-gauge needles to release their gonad primordium (consisting of 2 connected sister PGCs and 2 somatic gonad precursors). Germline-specific expression of GLH-1::GFP was used to identify PGCs, which were separated from gonad precursor cells by mouth pipetting using pulled glass capillaries coated with Sigmacote (Sigma, #SL2) and 1% BSA in egg buffer. 7.5 mg/mL pronase (Sigma, #P8811) and 5 mM EDTA were added to reduce sticking of gonad primordia to the poly-lysine coated slides and to weaken cell-cell interactions. Single sets of sister PGCs were transferred into 0.5 uL drops of egg buffer placed inside the caps of 0.5 mL low-bind tubes (USA Scientific, #1405-2600). Only PGCs that maintained bright fluorescence of GFP throughout isolation and were clearly separated from somatic gonad precursors were used for transcript profiling. Isolation of EGCs differed in 3 ways: 1) EGCs were dissected from L2 larvae that were fed for 20 hours after hatching, 2) the 2 EGCs that made up 1 sample may have come from different animals, and 3) the stage of each EGC could not be determined and therefore may have differed between samples. Tubes containing single sets of 2 sister PGCs or 2 EGCs were quickly centrifuged, flash frozen in liquid nitrogen, and stored at -80°C. A detailed protocol for isolating PGCs and EGCs from larvae is available upon request. At least 11 samples (replicates) of PGCs or EGCs were isolated for each condition.

### Isolation of RNAs from adult germlines

1st day hermaphrodite and male adult worms (approximately 20-24 hours post-mid-L4 stage) were cut open with 30-gauge needles in egg buffer (see recipe above, except not adjusted to 340 mOsm) containing 0.1% Tween and 1 mM levamisole to extrude their gonads. Gonads were cut at the narrow ‘bend’ to separate the gonad region containing mitotic and early meiotic germ cells from the region that contains oocytes and/or sperm; the former was used for RNA profiling. 20-60 gonads were mouth pipetted into 500 uL Trizol reagent (Life Technologies, #15596018), flash-frozen in liquid nitrogen, and stored at -80°C for up to 1 month before RNA extraction.

To release RNAs from gonads in Trizol, gonads were freeze-thawed 3x using liquid nitrogen and a 37°C water bath, while vortexing vigorously between cycles. RNAs immersed in Trizol were added to phase-lock heavy gel tubes (Brinkmann Instruments, INC., #955-15-404-5) and mixed with 100 uL of 1-bromo-3-chloropropane (BCP) (Sigma, #B9673), followed by room temperature incubation for 10 minutes. Samples were then centrifuged at 13,000g and 4°C for 15 minutes to separate phases. RNAs in the aqueous phase were precipitated by mixing well with 0.7-0.8x volumes of ice-cold isopropanol and 1 uL 20 mg/mL glycogen, followed by incubation at -80°C for 1-2 hours and centrifugation at 13,000g at 4°C for 30 minutes. RNA pellets were washed 3x with ice-cold 75% ethanol and then resuspended in 15 uL water. RNA concentration was determined by a Qubit fluorometer.

### Generation of cDNA sequencing libraries

*PGCs and EGCs:* Immediately after thawing PGCs and EGCs on ice, a 1:4,000,000 dilution of ERCC spike-in transcripts (Life Technologies, #4456740) was added to each sample. Double-stranded cDNAs from polyA(+) RNAs were generated using a SMART-seq method that combined parts of the Smart-seq2 (Picelli et al., 2014) and SMART-Seq v4 (Takara) protocols. Briefly, PGCs and EGCs were lysed at room temperature for 5 minutes in lysis buffer (Takara Bio, #635013) containing RNAse inhibitors (Takara Bio, #2313) to release mRNAs into solution. 1.2 µM custom DNA primer (IDT) was annealed to transcripts’ polyA tails by incubating samples at 72°C for 3 minutes and then immediately placing them on ice. Reverse transcription to generate double-stranded cDNA was performed using 200 U SmartScribe, 1x first-strand buffer, 2 mM DTT (Takara Bio, #639537), 1 mM dNTPs (Takara Bio, #639125), 4 mM MgCl_2_, 1 M betaine (Sigma, #B0300), 20 U RNAse Inhibitor (Takara Bio, #2313), and 1.2 µM custom template-switching oligo with a Locked Nucleic Acid analog (Qiagen) at 42°C for 90 minutes, followed by 70°C for 15 minutes to heat-inactivate the reverse transcriptase. cDNAs were PCR-amplified according to Takara’s SMART-Seq v4 protocol for 20 cycles using SeqAmp DNA Polymerase (Takara Bio, #638504) and a custom PCR primer. Amplified cDNAs were purified by SPRI using 1x Ampure XP beads (Agencourt, #A63881) and quantified using a Qubit fluorometer. All custom oligos contained a biotin group on their 5’ end to ameliorate oligo concatemerization. Illumina’s Nextera XT kit (Illumina, #FC-131-1096) was used with 350-400 pg cDNA as input to prepare dual-indexed Illumina RNA-sequencing libraries according to the manufacturer’s instructions. Libraries were PCR-amplified for 14 cycles and purified by SPRI using 0.6x Ampure XP beads.

#### Adult germlines

Illumina libraries were prepared from polyA(+) RNAs using the NEBNext Poly(A) mRNA Magnetic Isolation Module (NEB, #E7490) and the NEBNext Ultra RNA Library Prep Kit (NEB, #E7530) according to the manufacturer’s instructions. 100 ng total RNA was used to generate cDNAs, which were then amplified using 15 cycles of PCR.

Library concentration was measured by a Qubit fluorometer, and average fragment size was measured by a bioanalyzer or tapestation. Libraries were multiplexed and paired-end sequenced with 50 cycles on either an Illumina HiSeq2500 or NovaSeq 6000 SP flow cell at the QB3 Vincent J. Coates Genomics Sequencing Laboratory at UC Berkeley.

### Processing and analysis of RNA-seq data

*PGCs and EGCs:* Paired-end Illumina sequencing reads were aligned to the ce10 (WS220) genome downloaded from Ensembl using Hisat2 (2.2.1). Samtools (1.10) was used to remove duplicate reads and reads with low mapping quality (MAPQ < 10) from the alignment file. The function featureCounts from the subread package (2.0.1) (Liao et al., 2014) was used to obtain gene-level read (fragment) counts using a ce10 (WS220) transcriptome annotation file (Ensembl), which additionally contained 92 ERCC spike-in transcripts. 11 low-quality transcript profiles that were likely caused by a failure to capture mRNAs from 2 PGCs or EGCs, a well-known problem in single-cell RNA-sequencing, were identified using the R package scuttle (1.2.0) (McCarthy et al., 2017). For details of the transcript profiles generated in this study, including which were filtered out of our analysis, see our NCBI GEO accession (GSE198552). The R package DESeq2 (1.32.0) (Love et al., 2014) was used to perform Wald tests that identified differentially expressed genes in mutant vs wild-type samples. P-values were adjusted for multiple hypothesis testing by the Benjamini-Hochberg method to produce q-values. Genes with a q-value < 0.05 were considered differentially expressed. The scaling factors used by DESeq2 to normalize transcript profiles were calculated using the R package scran (1.20.1), which uses a pooling and deconvolution approach to deal with zero-inflation in low-input RNA-seq data (Lun et al., 2016). Bigwig files containing normalized read coverage over the WS220/ce10 genome were generated by bamCoverage from deepTools (Ramirez et al., 2016) using library scaling factors computed by scran, a bin size of 5, and a smoothing window of 15. Average read coverage across replicates was computed using WiggleTools (Zerbino et al., 2014) and visualized on the UCSC Genome Browser. Principal Component Analysis (PCA) was performed using DESeq2’s plotPCA function and variance-stabilized counts from DESeq2’s vst function. All visualizations of RNA-seq data were generated by the R packages ggplot2 (3.3.5) and ggpubr (0.4.0) or the UCSC Genome Browser.

#### Adult germlines

For transcript profiles from dissected adult germlines, data were processed as described above except the scaling normalization factors were computed by DESeq2.

### Gene sets

The germline-specific and germline-enriched gene sets were previously defined in Rechtsteiner et al. 2010 and Reinke et al. 2004, respectively. The soma-specific gene set was previously described in Knutson et al., 2017 and refined in this study by removing genes that have >5 TPM in our RNA-seq data from wild-type dissected adult germlines. The oogenesis and spermatogenesis gene sets were previously defined in Ortiz et al. 2014. Oogenesis genes are expressed at higher levels in dissected adult oogenic germlines than dissected adult spermatogenic germlines; spermatogenesis genes are the opposite.

### Single-molecule fluorescence in-situ hybridization (smFISH) in L1 larvae

100-200 gravid adult mothers were allowed to lay offspring in drops of S basal overnight. Starvation-synchronized L1 offspring were collected and fed HB101 bacteria in S basal for 5 hours. Fed L1s were washed 3-4 times with S basal to remove bacteria and then used for smFISH using the protocol described in Ji and van Oudernaarden, 2012 with a few modifications. Briefly, L1s were fixed with 3.7% formaldehyde for 45 minutes at room temperature, followed by 3 washes with PBS-Tween (0.1%). Fixed L1s were incubated in 75% ethanol at 4°C overnight and up to 3 days. RNAs were hybridized to 25 nM RNA probe sets in hybridization buffer (2x SSC, 10% formamide, 0.1% Tween-20, and 0.1 g/mL dextran sulfate) at 37°C overnight. Afterwards, larvae were washed 2x in hybridization buffer at 37°C for 30 minutes, the second of which included 1 ng/uL of DAPI. Larvae were washed 3x in PBS-Tween (0.1%) and mounted in anti-fade medium consisting of n-propyl gallate and Vectashield (Vector Laboratories, #H-1000). Mounted samples were immediately imaged on a spinning disk confocal microscope using a 100x oil objective to acquire 3D Z-stacks of PGCs; only Z-slices containing GLH-1::GFP signal were imaged. All RNA probe sets were conjugated to Quasar 670 fluorescent dye. We designed and purchased the *lsd-1*, *mbk-1*, *pek-1*, and *lin-15B* probe sets from Stellaris. The *cpg-2, pgl-3,* and *chs-1* probe sets were gifts from Dr. Erin Osborne Nishimura.

### Counting transcripts in 3D smFISH images

To batch process raw smFISH 3D images into transcript abundance measurements in PGCs, a custom pipeline written in Fiji (v2.1.0/1.53C) and MATLAB (R2020a) was developed. Much of the MATLAB code and strategy was adapted from Raj et al. 2008 to create our pipeline. Images were removed from downstream analyses if they were obviously very dim in all channels. Numbers of analyzed PGCs per probe set per genotype are indicated in Figure 2. A Laplacian of Gaussian (LoG) filter was used to enhance the signal-to-noise contrast in smFISH images. Those filtered 3D images were thresholded by signal intensity to produce binary images. The ‘imregionalmax’ function from the Imaging Processing Toolbox in MATLAB was used to find and count regional maxima (transcript foci) in those binary images. To isolate and count regional maxima in PGCs, a 2D binary image mask of the PGCs was generated in Fiji from a maximum-intensity Z-projection of the GLH-1::GFP image channel and applied to all Z-slices in the 3D smFISH images. Segmentation of PGCs to create the 2D mask was done in Fiji by first blurring Z-projections using a large Gaussian kernel and then detecting edges in the blurred image. Mann-Whitney tests were performed to compare transcript abundance between *mes-4* and wild-type PGCs.

To choose an appropriate signal intensity threshold for a 3D smFISH image, the number of detected regional maxima across 100 increasingly stringent thresholds applied to the image were plotted (Raj et al., 2008). In that plot, a range of thresholds that produced similar numbers of detected regional maxima was identified, and a threshold within that range was selected. Since threshold values were similar for all images within an image set (the collection of images acquired on the same day and for one probe set), an averaged threshold (across 5 images) was calculated and applied to all images in the set. Notably, threshold values were similar across different image sets. Manual counting of dots in a few smFISH images gave similar values as our semi-automated pipeline.

### Germline size analysis

All analyses were performed by live imaging 1st day adults (approximately 20-24 hours post-mid-L4 stage) and evaluating germline size using the germline marker GLH-1::GFP. Adult germlines were classified into 1 of 3 categories: (1) ‘Full’ if its size was similar to that of a wild-type adult germline, (2) ‘Partial’ if it had at least ∼15 GLH-1::GFP(+) germ cells but wasn’t large enough to be classified as ‘full’, and (3) ‘Absent/tiny’ if it had < 15 GLH-1::GFP(+) germ cells. In rare ambiguous cases where a germline’s size was intermediate between categories, the germline was assigned to the category of the smaller size. To classify germline size for hermaphrodites, which have 2 gonad arms, only the size of the larger gonad arm was considered; in most cases, both gonad arms were similar in size. 2-tailed Fisher’s exact tests were performed to test whether the proportion of worms with a full germline is higher or lower in 1 sample vs another sample.

Confocal microscopy was used to acquire live images of scored germlines in adults. Live adults were placed in a drop of 1 uL H_2_O and 1 uL polystyrene microspheres (Polysciences, Inc., #00876) and then immobilized on 6% agarose pads. Images were acquired in Z-stacks using a 20x air objective and then converted to Z-projections of maximum intensity using Fiji. DIC projections were used to outline the body of worms.

### Gamete and progeny analysis

To determine whether *mes-4* M-Z- X^sp^ males that made full-sized germlines also produced sperm, we imaged ethanol-fixed and DAPI-stained males using a spinning disk confocal microscope. Males were assigned to 1 of 3 bins by the number of sperm (small, dense DAPI-stained bodies in the proximal region of the germline) they contained: 1) 0-10 sperm, 2) 10-50 sperm, or 3) >50 sperm. To determine whether *mes-4* M-Z- X^sp^/X^sp^ hermaphrodites that made full-sized germlines also produced progeny, live-imaging was used to determine if they contained at least 1 egg in their uterus, after which single egg-containing hermaphrodites were transferred to individual plates and scored for production of viable progeny. In this way, hermaphrodites were assigned to 1 of 3 bins: 1) ‘no eggs’, 2) ‘eggs’ (but did not produce viable progeny, and 3) ‘eggs and progeny’. Only males and hermaphrodites with full-sized germlines were considered in the analysis. We used 2-sided Fisher’s exact tests to compare what proportion of wild-type and *mes-4* M-Z- mutant males contained >50 sperm and hermaphrodites contained ‘eggs and progeny’. P-value designation is ** < 0.001.

### Tracking X-chromosome inheritance patterns

Methods used to track X-chromosome inheritance are diagrammed in Figure 3—figure supplement 1. To identify F1 male offspring that received their single X from the oocyte (X^oo^) or the sperm (X^sp^), crosses were performed with 1 parent contributing an X-linked *eft-3p::mCherry* transgene. F1 male offspring that inherited the transgene were easily distinguished by bright cytoplasmic mCherry fluorescence in their soma. A *gpr-1* over-expression allele was used to generate F1 hermaphrodite offspring whose germline inherited either 2 genomes from the sperm or 2 genomes from the oocyte (Besseling and Bringmann, 2016; Artiles et al., 2019). Those non-Mendelian offspring were visually identified by patterns of mCherry fluorescence in their pharyngeal muscle cells (*myo-2p::mCherry* X) (Figure 3—figure supplement 1).

### RNA interference (RNAi) depletion of gene products

RNAi was performed by feeding worms *E. coli* HT115 bacteria that carry a gene target’s DNA sequence in the L4440 vector (Kamath and Ahringer, 2003). Most RNAi constructs were obtained from the Ahringer RNAi library and sequence confirmed. *lin-15B*, *lsy-2*, *nfya-1*, *eor-1*, and *sma-9* RNAi constructs were generated for this study. RNAi constructs were streaked onto LB agar plates containing 100 ug/mL carbenicillin and 10 ug/mL tetracycline. Single clones were cultured overnight (14-17 hours) at 37°C in LB and carbenicillin (100 ug/mL). The following day, RNAi cultures were spotted onto 6-cm NGM plates containing 1 mM IPTG and 100 ug/mL carbenicillin (both added by top spreading), and then left to dry for 2 days at room temperature in the dark. To deplete both maternal load and embryo-synthesized gene product, we placed L4-stage larval mothers onto RNAi plates, grew them 1 day to reach adulthood, then moved those adults to new RNAi plates to lay offspring. RNAi was done at 20°C. Germline size was scored in 1st day adult offspring as described above.

### Statistical analysis

Sample sizes for worms scored for germline size and for images used to count transcripts in smFISH analysis are indicated in the respective figures. Gene set enrichment analyses were performed using hypergeometric tests in R and the DAVID Bioinformatics Resource 6.8 (Huang et al., 2009). The sizes of gene sets used for those tests are noted in the respective figure legends. P-value designations are * < 0.01, ** < .001, *** < 1e-4, and **** < 1e-5.

### Spinning-disk confocal microscopy

All images were acquired using a spinning-disk confocal microscope equipped with a Yokogawa CSU-X1 confocal scanner unit, Nikon TE2000-E inverted stand, Hamamatsu ImageEM X2-CCD camera, solid state 405, 488, 561, 640 nm laser lines, 460/50, 525/50, 593/40, 700/75 nm (EM/BP) fluorescence filters, DIC, Nikon Plan Apo VC 20x/0.5 air objective, Nikon Plan Apo 100x/1.40 oil objective, and Micro-Manager software (1.4.20). Image processing for images was done in Fiji (2.1.0/1.53C) and photoshop.

### Transcription factor analyses

#### Analyses of ChIP data

Bed files containing transcription factor (TF) binding sites (ChIP-chip or ChIP-seq ‘peaks’) across the ce10 genome were downloaded from the modENCODE (Gerstein et al., 2010) and modERN (Kudron et al., 2018) websites. Each bed file was loaded into R and converted into a GRanges object using the GenomicRanges package (Lawrence et al., 2013). TF binding sites were assigned to genes using the package ChIPSeeker (Yu et al., 2015). A TF binding site was assigned to a gene if it overlapped with a gene’s TSS region (500 bp upstream of TSS). If a TF had more than 1 set of binding site data (e.g. 2 ChIP-seq experiments), each set was processed separately and then the TF-assigned genes were merged. TFs enriched in promoters of X-linked UP genes in *mes-4* PGCs and/oror EGCs (584 genes) compared to all X-linked genes (2808) were identified using hypergeometric tests (p-value < 0.05).

#### Analyses of DNA motifs

Position-weight matrices for known *C. elegans* TF DNA motifs were downloaded from the CisBP database (Weirauch et al., 2014). X-linked genes that contain a TF’s motif in their promoter (500 bp upstream of TSS) were identified using FIMO from the Meme suite in R using default parameters (Grant et al., 2011). Motifs significantly enriched in promoters of X-linked UP genes in *mes-4* PGCs or EGCs (584 genes) compared to all X-linked genes (2808) were identified using hypergeometric tests (p-value < 0.05).

## DATA AVAILABILITY

All sequencing data generated in this study were deposited in NCBI GEO under accession GSE198552.

## ACKNOWLEDGEMENTS

We thank past and present members of the Strome lab and Grant Hartzog for helpful discussions, Ben Abrams for microscopy advice, and Joshua Arribere for help with statistical analyses. We thank Erin Osborne Nishimura for kindly gifting RNA probe sets for smFISH. This work was supported by NIH grants T32GM008646 and F31GM125305 to C.C. and R01GM34059 to S.S.

## SUPPLEMENT

**Figure 1—figure supplement 1:**
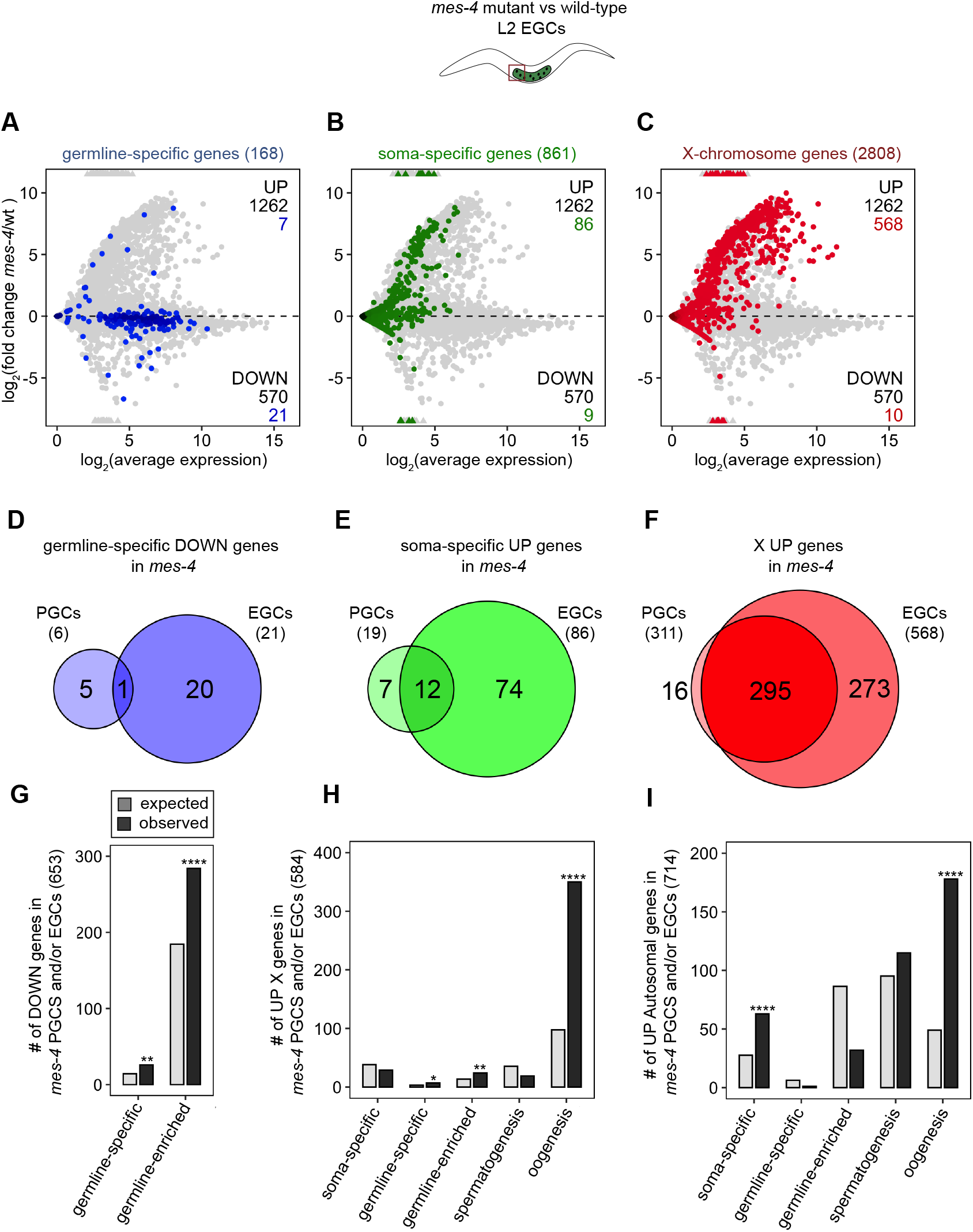
*mes-4* M-Z- EGCs have more severe transcriptome defects than *mes-4* M-Z- PGCs. **(A,B,C)** MA plots as described in Figure 1B-D showing differential expression analysis for *mes-4* vs wt Early Germ Cells (EGCs). (D,E,F) Venn diagrams comparing sets of mis-regulated genes (*mes-4* vs wt) between PGCs and EGCs: (D) DOWN germline-specific genes, (E) UP soma-specific genes, and (F) UP X genes. (G,H,I) Bar plots showing the expected number (light gray) and observed number (dark gray) of mis-regulated genes in either *mes-4* PGCs and/or EGCs that are members of the indicated gene sets. Hypergeometric tests were performed in R to test for gene-set enrichment. P-value designations are * < 0.01, ** < 0.001, and **** < 1e-5. (G) Enrichment analyses for DOWN genes were restricted to 6,682 protein- coding genes that we defined as ‘expressed’ (minimum average read count of 1) in either wt PGCs or EGCs. Gene set sizes germline-specific (146), germline-enriched (1887). (H,I) Enrichment analyses for UP genes included all 20,258 protein-coding genes in the transcriptome. Gene set sizes (on the X and autosomes):: soma-specific (184, 677), germline-specific (15, 153), germline-enriched (65, 2111), spermatogenesis (171, 2327), oogenesis (470, 1201).

**Figure 1—figure supplement 2:**
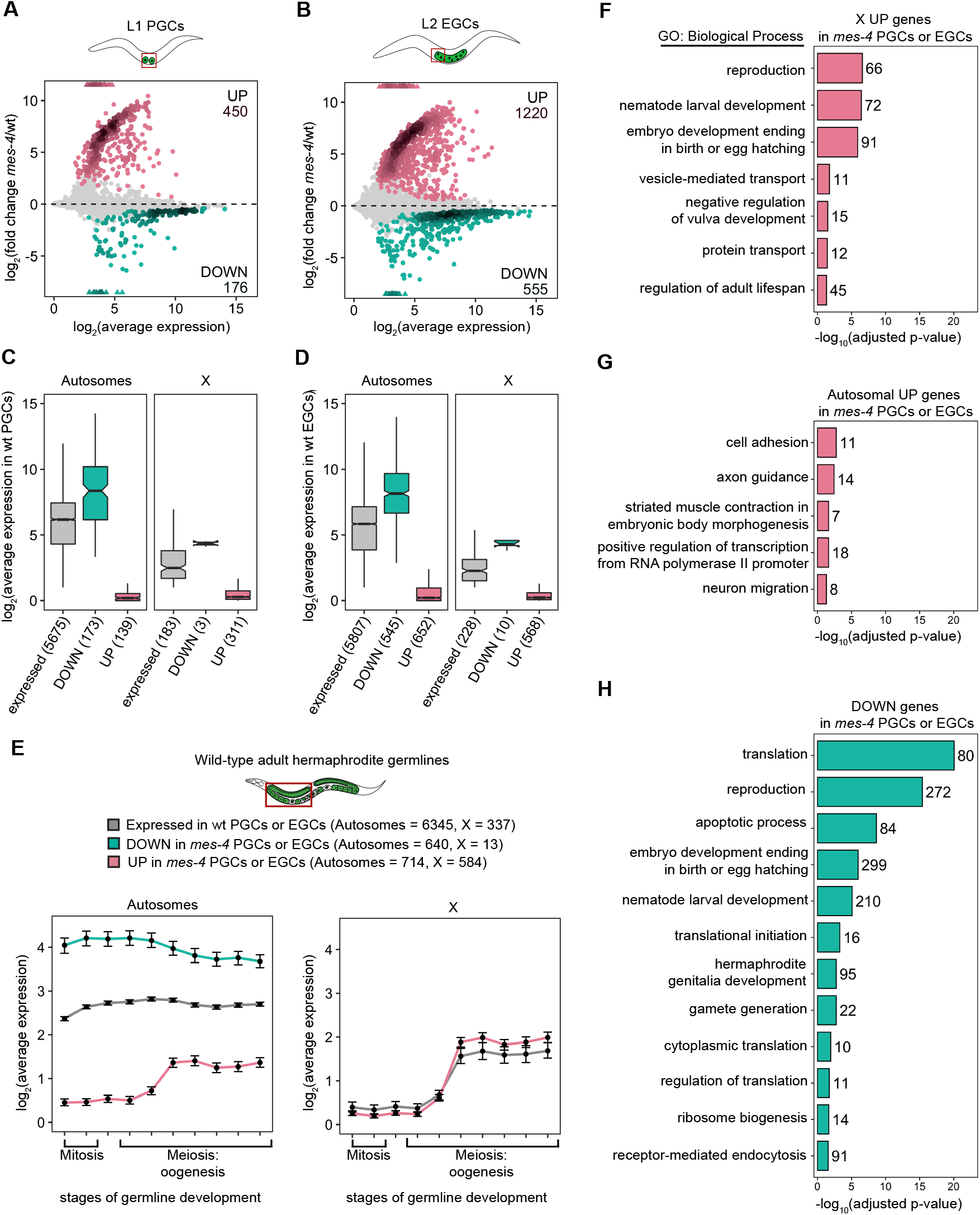
Analysis of features of mis-regulated genes in *mes-4* M-Z- PGCs and EGCs. (A,B) MA plots as described in Figure 1B-D showing differential expression analysis for (A) *mes-4* vs wt PGCs and (B) *mes-4* vs wt EGCs. Colored circles represent mis-regulated genes (q-value < 0.05); UP genes are in pink and DOWN genes are in teal. (C,D) Boxplots showing distributions of transcript abundance for autosomal (left) or X-linked (right) expressed genes (gray), DOWN genes (teal), UP genes (pink) in (C) wt PGCs and (D) wt EGCs. Boxplots show the median, the 25^th^ and 50^th^ percentiles (boxes), and the 2.5^th^ and 97.5^th^ percentiles (whiskers). (E) Line plots showing log_2_(average expression) across 10 regions of a wild-type adult hermaphrodite gonad that capture different stages of germline development (data from Tzur et al., 2018, Tables S2A and S10A). Average expression values were calculated in Tzur et al. by normalizing each gene’s read counts to the total number of read counts in a sample, multiplying those normalized values by 10^5^, and finally averaging across 2 independent samples. Each dot represents the mean expression of a gene set (colors) in 1 of the 10 germline regions. Whiskers correspond to 95% confidence intervals of the mean. Gene set sizes are indicated. The set of DOWN X genes was not analyzed due to its small size (13 genes). (F,G,H) Bar plots showing significantly enriched (Benjamini Hochberg-adjusted p-value < 0.05) gene ontology (GO) biological process terms in sets of mis-regulated genes (*mes-4* PGCs and/or EGCs): (F) X UP genes, (G) autosomal UP genes, and (H) DOWN genes. Numbers of mis-regulated genes for each GO term are indicated. Analyses of UP genes (F and G) used all 20,258 protein coding genes as a background, while analysis of DOWN genes (H) used our set of 6,345 protein-coding genes expressed in wt PGCs or EGCs as a background. All GO analyses were performed using DAVID Bioinformatics Resource 6.8 (Huang et al., 2009).

**Figure 2—figure supplement 1:**
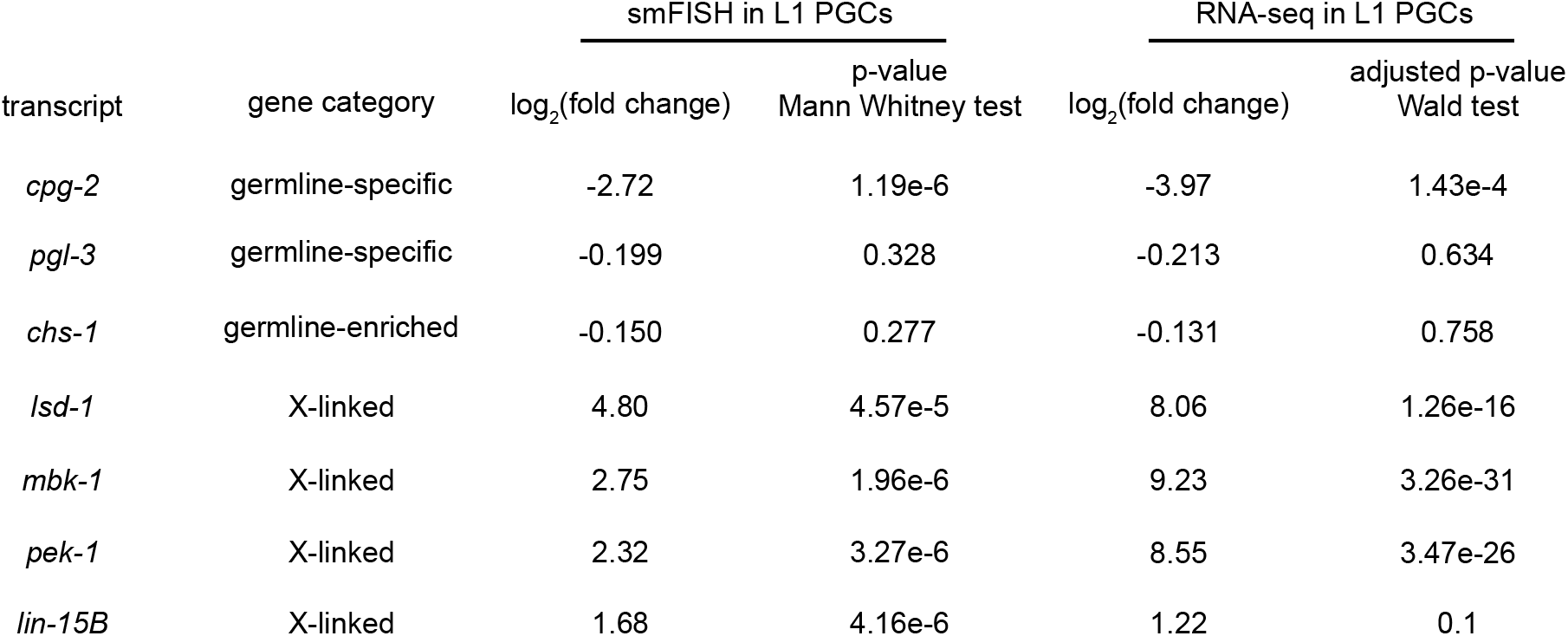
Comparison of smFISH and transcript profiling data. Table showing differential expression statistics for the 7 genes tested by smFISH. Log_2_(fold change) values were calculated by comparing mean transcript counts in *mes-4* PGCs to wt PGCs, and the p-value is from a Mann Whitney test. For transcript profiling analysis, ‘shrunken’ log_2_(fold change) values were calculated using DESeq2 and ashr in R (Stephens, 2017).

**Figure 3—figure supplement 1:**
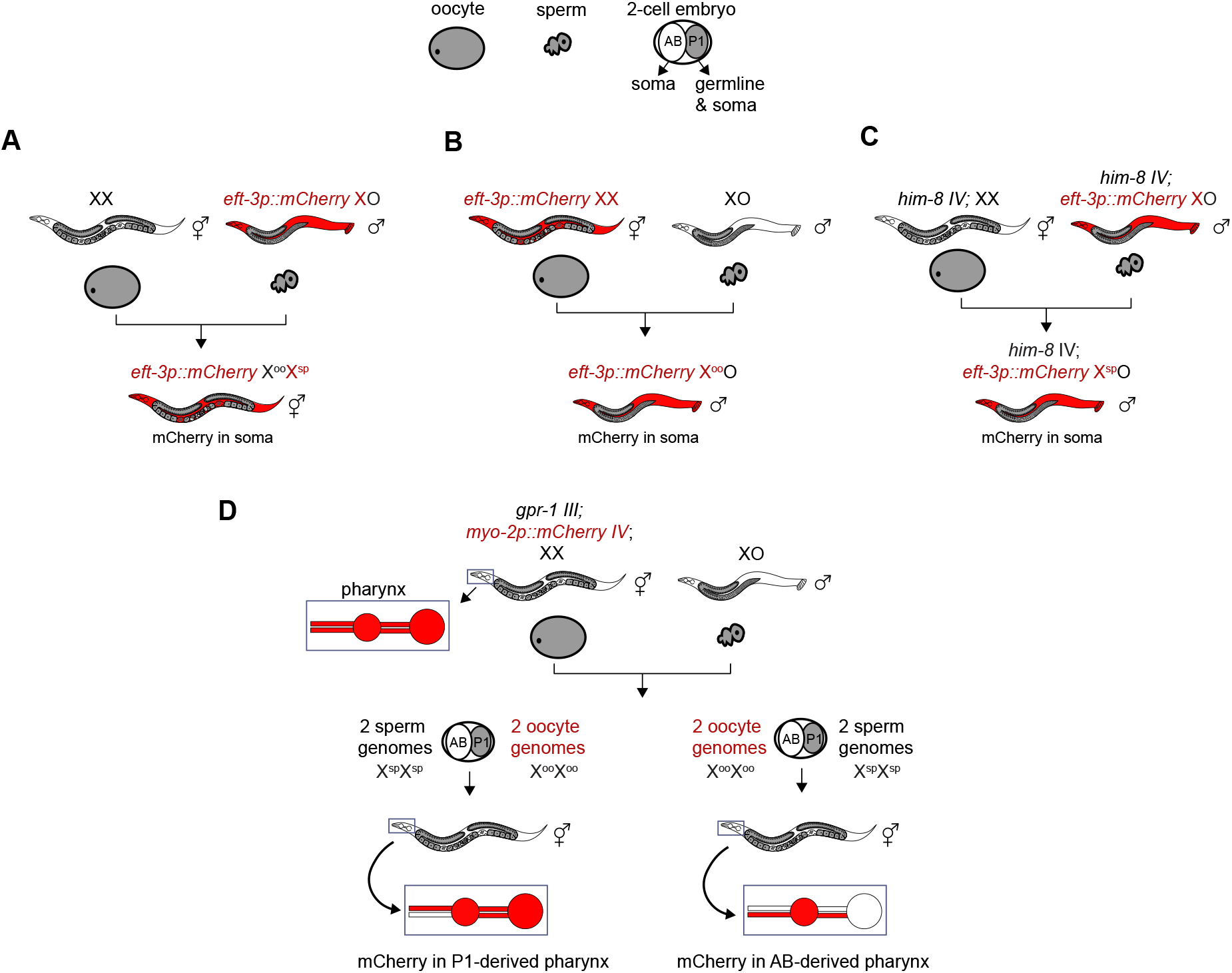
Genetic strategies to generate and identify F1 offspring that inherited different X-chromosome endowments from parents. (A-C) X-linked and soma-expressed mCherry driven by the *eft-3* promoter was used to track X-chromosome inheritance patterns in F1 offspring. (C) To generate males that inherited their single X from the sperm, we used the *him-8(e1489)* allele to cause X-chromosome nondisjunction in the maternal germline, which causes some oocytes to lack an X. (D) To generate hermaphrodites whose germline inherited either 2 genomes from the oocyte or 2 genomes from the sperm, we used a *gpr-1* over-expression allele. Expression of mCherry driven by the *myo-2* promoter in AB-derived pharyngeal muscle or P1-derived pharyngeal muscle was used to identify F1 hermaphrodite offspring with non-Mendelian inheritance patterns.

**Figure 3—figure supplement 2:**
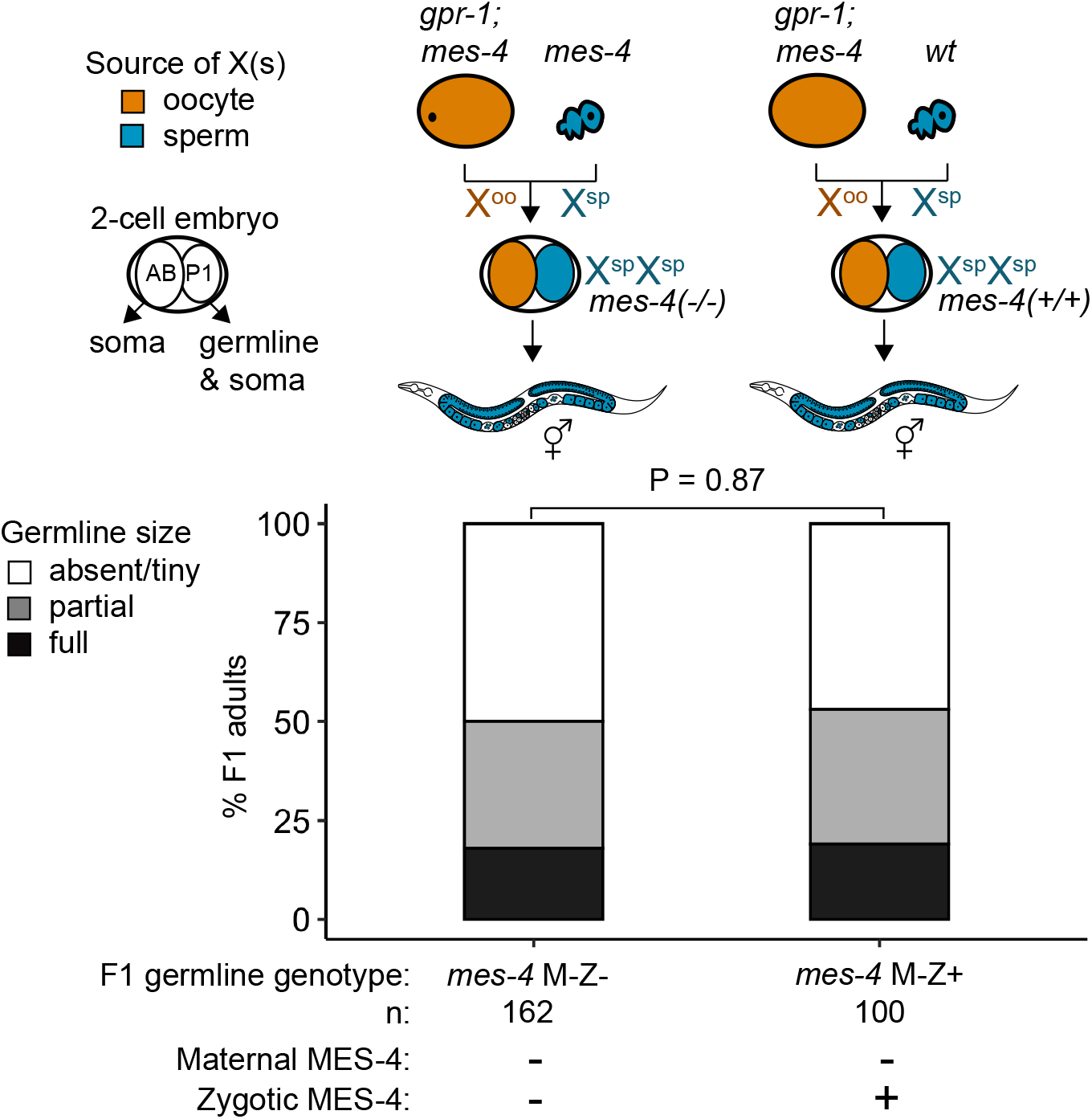
*mes-4* M-Z+ X^sp^/X^sp^ mutants do not have healthier germlines than *mes-4* M-Z- X^sp^/X^sp^ mutants. Bar plots as described in Figure 3 showing distributions of germline size (absent/tiny, partial, and full) in F1 offspring whose germline inherited 2 X chromosomes from the sperm. Numbers of scored F1 offspring, the genotype of their germlines, and the presence or absence of maternal MES-4 or zygotic MES-4 are indicated below the bars. Notably, the germlines of *mes-4* M-Z+ offspring have 2 wild-type copies of *mes-4* that they inherited from the sperm.

**Figure 3—figure supplement 3:**
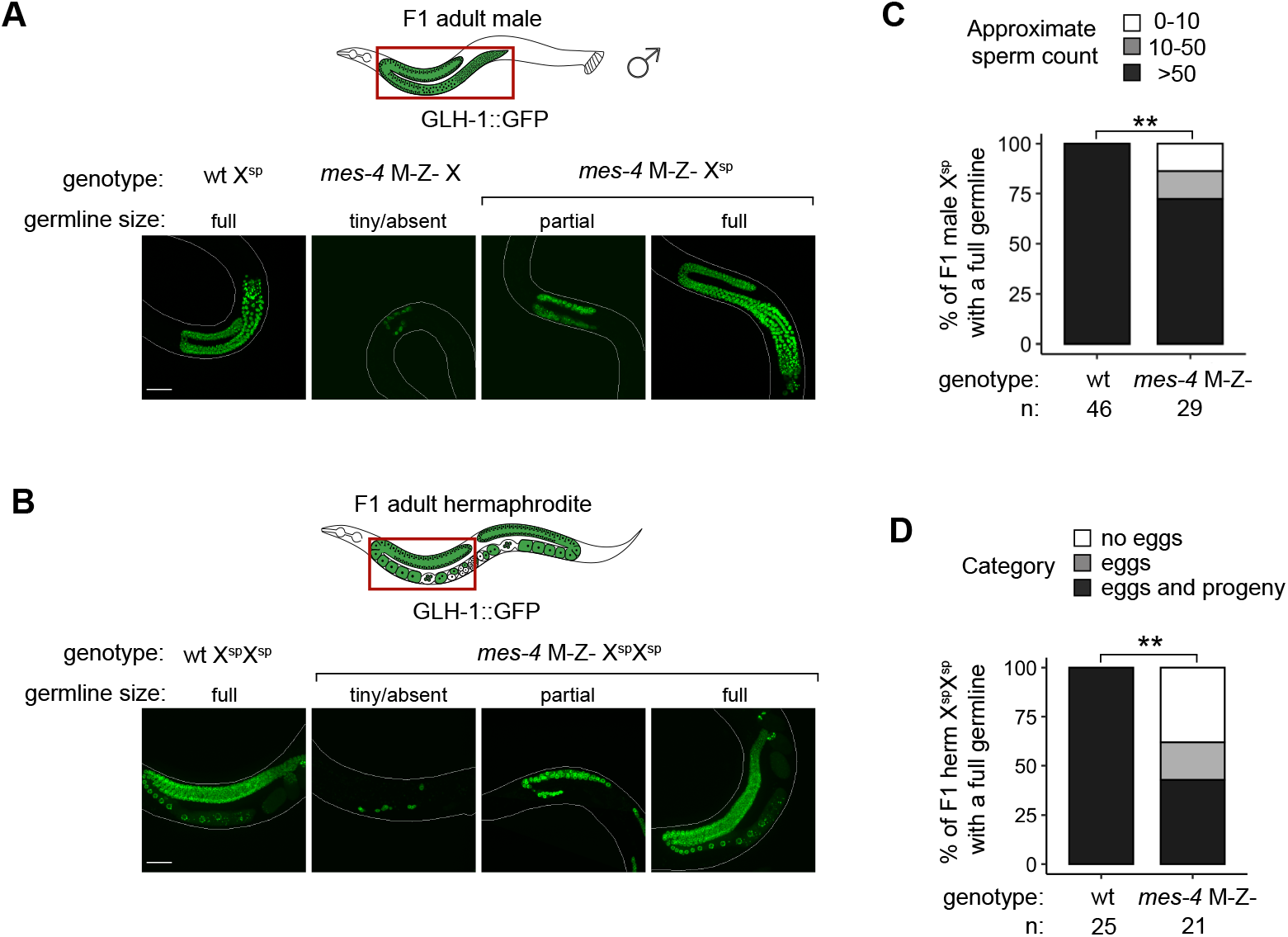
Further fertility analyses of *mes-4* M-Z- mutants that inherited their X(s) from the sperm. (A,B) Representative images of live F1 offspring with a germline scored as either tiny/absent, partial, or full. Genotypes of imaged worms with respect to the *mes-4* locus and their germline’s X-chromosome composition are indicated. Green signal is the germline marker GLH-1::GFP. Worm bodies are outlined in white. Scale bar is 45 µM. (A) F1 male offspring generated using the *him-8(e1489)* allele. All imaged males inherited their single X from the sperm, except the male with a tiny/absent germline; that male inherited its X either from the mother’s oocyte or mother’s sperm (self-fertilization). (B) F1 hermaphrodite offspring generated using the *gpr-1* allele form a germline composed of 2 genomes from the sperm. (C) Bar plots showing sperm counts in F1 male offspring that were classified as having a full germline. Percentages of F1 males that had >50 sperm were compared between wt and *mes-4* mutant populations using a 2-sided Fisher’s exact test. P-value designation is ** < 0.001. (D) Bar plots showing the presence or absence of eggs and viable progeny in F1 hermaphrodite offspring that were classified as having full germlines. Percentages of F1 hermaphrodites that had eggs and viable progeny were compared between wt and *mes-4* mutant populations using a 2-sided Fisher’s exact test.

**Figure 4—figure supplement 1:**
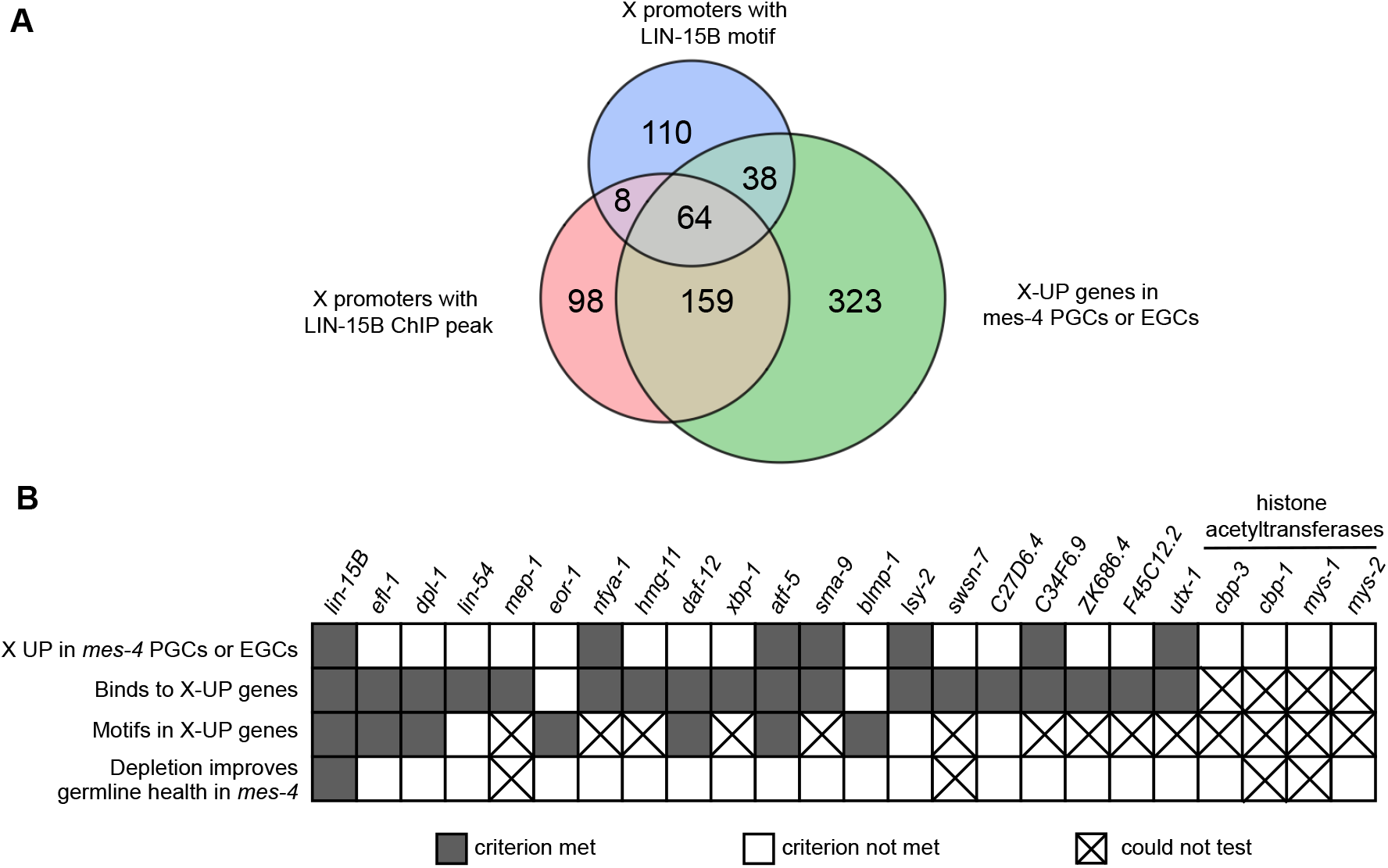
Identification and testing of candidate transcription factors for a role in causing sterility of *mes-4* M-Z- mutants. (A) Venn diagram comparing 3 gene sets: X UP genes in *mes-4* PGCs and/or EGCs, X genes with a LIN-15B ChIP peak in their promoter region (500 bp upstream of TSS), and X genes with a LIN-15B binding motif in their promoter region. LIN-15B ChIP-chip data were from embryos (downloaded from modENCODE), and LIN- 15B ChIP-seq data from L3 land L4 larvae (downloaded from modERN). (B) Table showing the candidate transcription factors that were tested for a role in causing sterility of *mes-4* M-Z- worms (columns) and the criteria used to select them (rows). Gray-colored squares indicate a criterion was met, white-colored squares indicate a criterion was not met, and an ‘X’ indicates a criterion could not be tested or the effect of RNAi on germline size could not be determined. RNAi of *cbp-1, mys-1*, or *swsn-7* caused highly penetrant embryo and larval lethality, which prevented us from sampling a sufficient size of healthy adults. RNAi of *mep-1* transcripts caused ectopic expression of GLH-1::GFP(+) P-granules in somatic tissues, which complicated our identification of germline tissue by live imaging GLH-1::GFP. The RNAi phenotypes we observed for *cbp-1*, *swsn-7*, and *mep-1* have been previously reported (Maeda et al., 2001; Unhavaithaya et al., 2002; Cabianca et al., 2019).

**Figure 4—figure supplement 2:**
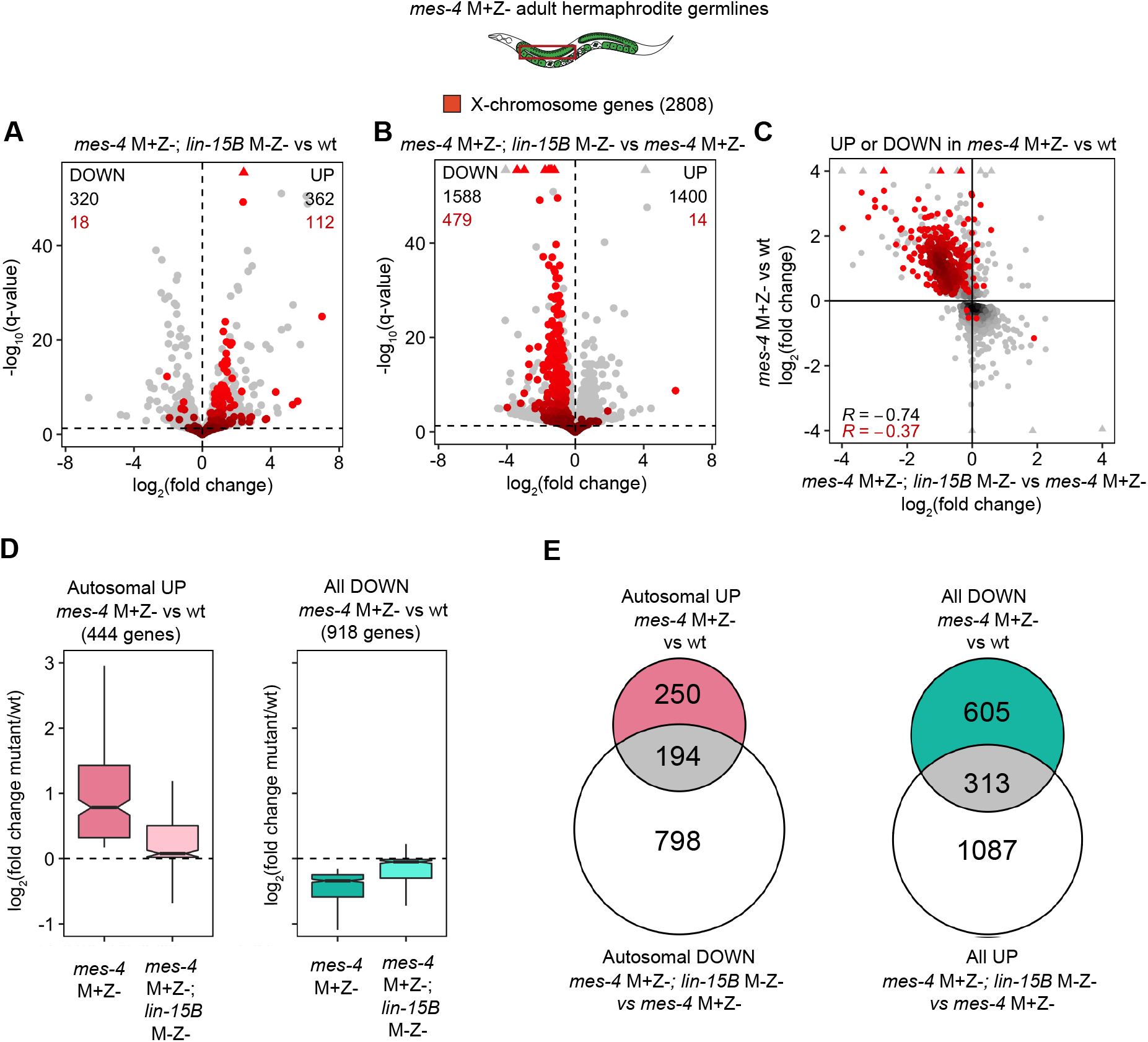
Further analysis of how LIN-15B impacts the transcriptome of *mes-4* M+Z- dissected adult germlines. (A,B) Volcano plots as described in Figure 4A showing differential expression analysis for (A) *mes-4* M+Z-; *lin-15B* M-Z- vs wt adult germlines and for (B) *mes-4* M+Z-; *lin-15B* M-Z- vs *mes-4* M+Z- germlines. X genes are colored red. (C) Scatterplot comparing log_2_(fold change) in *mes-4* M+Z- vs wt adult germlines to log_2_(fold change) in *mes-4* M+Z-; *lin-15B* M-Z- vs *mes-4* M+Z- adult germlines. Only genes that were either UP or DOWN (q-value < 0.05) in *mes-4* M+Z- vs wt adult germlines are shown. X genes are colored red. The Spearman’s correlation coefficient is indicated for all plotted genes (black) and for plotted X genes (red). (D) Boxplots as described in Figure 4B for autosomal UP genes in *mes-4* M+Z- vs wt (left) and for DOWN genes in *mes-4* M+Z- vs wt (right). (E) Venn diagrams as described in Figure 4C for the indicated gene sets.

**Figure 4—figure supplement 3:**
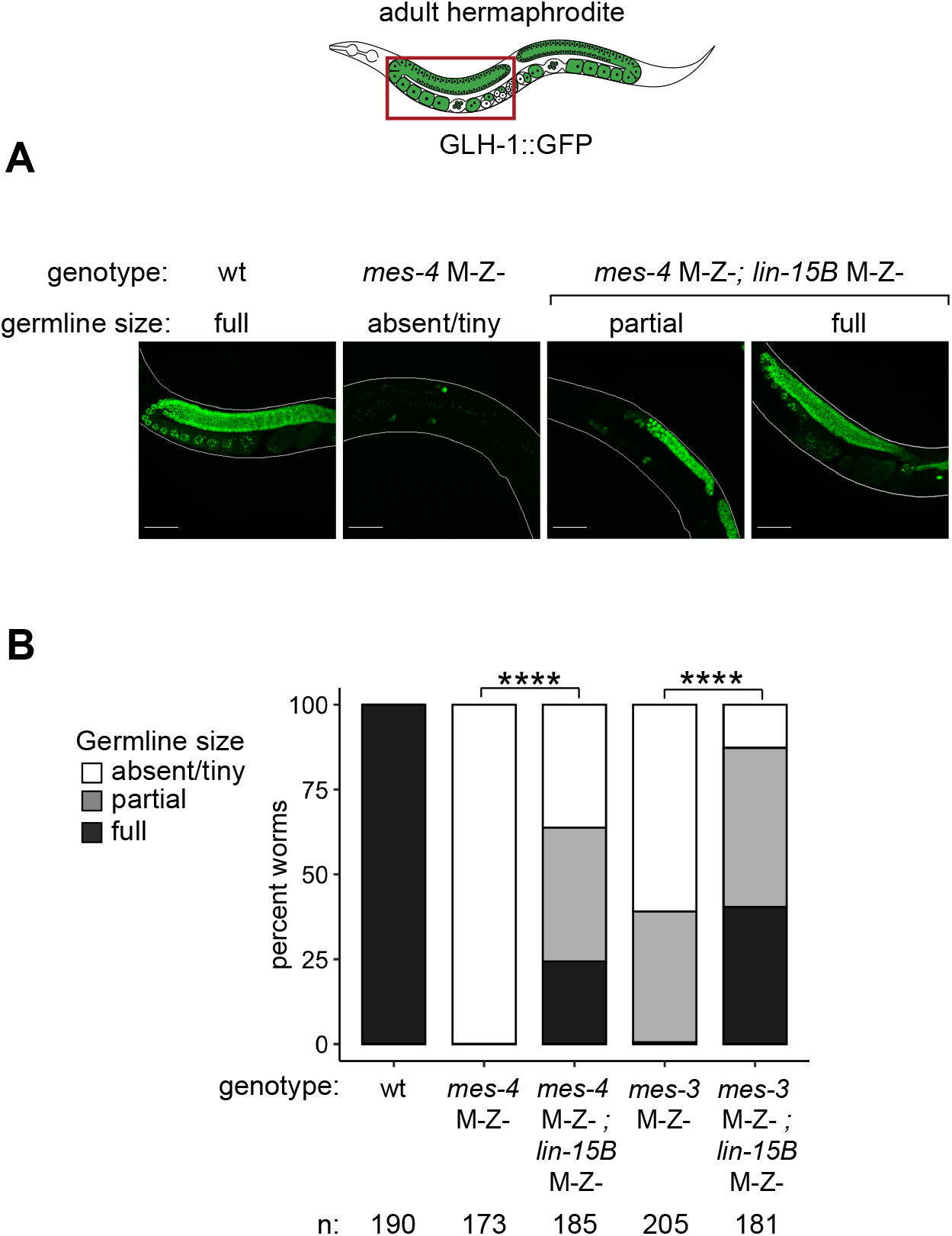
Removal of LIN-15B improves germline health in *mes-4* M-Z- X^oo^/X^sp^ mutant hermaphrodites. (A) Representative images of live F1 adult hermaphrodite offspring with a germline scored as either absent/tiny, partial, or full. Genotypes of imaged worms with respect to *mes-4* and *lin-15B* are indicated. All F1s inherited 1 X from the oocyte and 1 X from the sperm. Green signal is the germline marker GLH-1::GFP. Worm bodies are outlined in white. Scale bar is 45 µM. (B) Bar plots as described in Figure 4D showing distributions of germline size in F1 offspring. Genotypes and sample sizes are indicated. Percentages of F1s with full-sized germlines were compared using 2-sided Fisher’s exact tests. P-value designation is **** < 1e-5.

**Figure 5—figure supplement 1:**
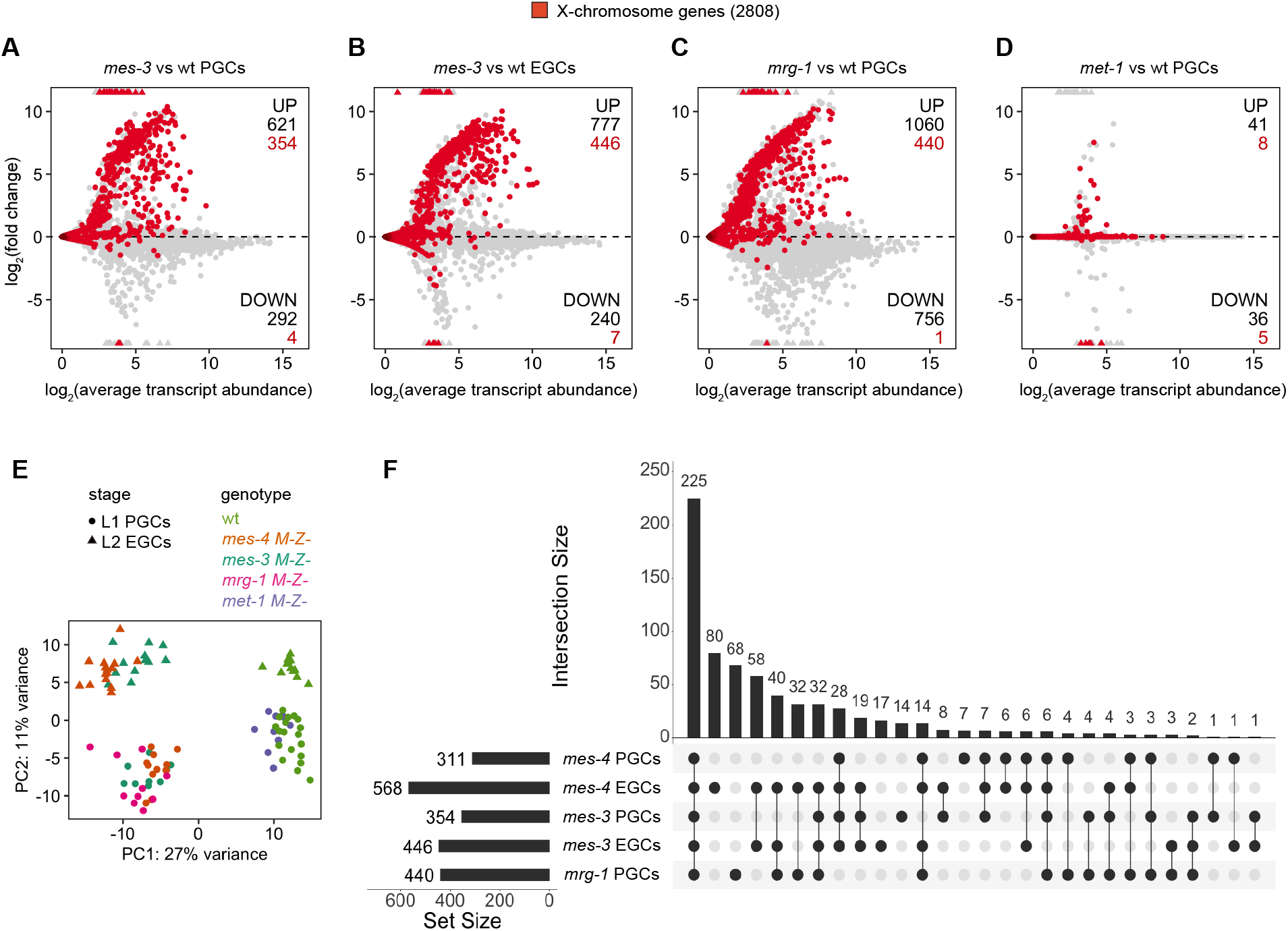
Comparison of X mis-expression in PGCs and EGCs dissected from various chromatin regulator mutants. (A-D) MA plots as described in Figure 1B-D showing differential expression analysis for (A) *mes-3* M-Z- vs wt PGCs, (B) *mes-3* M-Z- vs wt EGCs, (C) *mrg-1* M-Z- vs wt PGCs, and (D) *met-1 M-Z-* vs wt PGCs. X genes are colored red. (E) PCA plot showing all analyzed transcript profiles across the first 2 principal components. Variance-stabilized counts computed by the vst function in DESeq2 were used to perform PCA. Stages and genotypes of worms are indicated by shape and color, respectively. (F) Upset plot generated using the UpsetR R package comparing sets of X UP (mutant vs wt) genes across *mes-4* PGCs, *mes-4* EGCs, *mes-3* PGCs, *mes-3* EGCs, and *mrg-1* PGCs.

